# The offensive role of the *Bacillus* extracellular matrix in driving metabolite-mediated dialogue and adaptive strategies with pathogenic fungi

**DOI:** 10.1101/2025.02.06.636830

**Authors:** Alicia I. Pérez-Lorente, Carlos Molina-Santiago, David Vela-Corcía, Paolo Stincone, Jesús Hierrezuelo, Montserrat Grifé, Abzer K. Pakkir Shah, Antonio de Vicente, Daniel Petras, Diego Romero

## Abstract

Bacterial□fungal interactions have traditionally been attributed to secondary metabolites, but the role of the bacterial extracellular matrix (ECM) in shaping these relationships has remained unclear. Here, we demonstrate that the ECM protein TasA is a key mediator in the antagonistic interaction between *Bacillus subtilis* and *Botrytis cinerea*. TasA enables *Bacillus* to tightly adhere to fungal hyphae, disrupts the β-glucan layer, and compromises fungal cytoskeletal integrity synergistically with fengycin, which causes cytological damage. Additionally, TasA acts as a carrier for bacillaene, amplifying its fungistatic activity. In response, *B. cinerea* mounts a multifaceted defense, enzymatically degrading fengycin, producing antibacterial oxylipins, and activating adaptive programs such as hyphal branching and chlamydospore formation. Our findings reveal the previously unrecognized role of ECM components in fungal suppression and the modulation of fungal adaptive responses. This study reveals the complex interplay between microbial aggression and defense, providing new insights into the ecological dynamics of microbial competition and coexistence.

## Introduction

Microbial interactions shape microbial community dynamics and impact ecosystems, particularly within the plant microbiome^1–3^. These interactions range from cooperative to competitive and are often mediated by secondary metabolites, signaling molecules, and biofilms that influence plant health^4^. Beneficial microbes support nutrient acquisition, plant defense, and stress tolerance but compete for limited resources^5,6^ with detrimental pathogens, including fungal species such as *Botrytis cinerea* ^7–10^.

*Bacillus subtilis* is a gram-positive bacterium known for its ability to promote plant growth and effectively antagonize fungal pathogens such as *B. cinerea*, the causative agent of gray mold disease, which significantly impacts crops and leads to economic losses ^7–10^. The antagonistic activity of *B. subtilis* stems from its production of antifungal lipopeptides, such as fengycin and surfactin, and apparently robust biofilm formation capacity^11^. The extracellular matrix (ECM) of *B. subtilis* comprises mainly but not exclusively exopolysaccharides (EPSs), TasA, TapA, and BslA^12,13,14^, which provide structural integrity and participate in interactions with plants; bacteria such as *Pseudomonas* species^15,16^; and fungal species such as *Aspergillus niger* ^17,18^ or *Rhizophagus irregularis* ^19^. However, the exact role of the ECM in interactions with phytopathogenic fungi remains poorly understood.

Traditionally, the *B. subtilis–B. cinerea* interaction has been viewed as being statically antagonistic. However, microbial interactions are dynamic, with both organisms undergoing metabolic and transcriptomic changes^20^. While *B. subtilis* suppresses *B. cinerea* growth, a portion of the fungal population adapts and survives^21–23^. This study demonstrates the multifaceted involvement of the ECM, beyond adhesion, in this interkingdom chemical crosstalk and in antagonistic interactions, with TasA and fengycin as key components driving fungal modulation and ecological balance within microbial ecosystems.

## Results

### The *Bacillus* ECM mediates adhesion and bidirectional damage in the interaction with *Botrytis*

Microbial interactions are inherently dynamic and are shaped by evolving exchanges between organisms and their environment. This ecological fluidity is especially pronounced in antagonistic or competitive relationships within microbial communities, where organisms constantly adapt to maintain ecological balance or eliminate competitors^24–27^. Historically, the interaction between *B. subtilis* and *B. cinerea* has been described as unidirectionally antagonistic, driven by the ability of *B. subtilis* to suppress fungal growth through the secretion of antifungal compounds, including plipastatins and surfactins^28–30^. However, our observations revealed a more complex and dynamic interaction over time. In the short term, *Bacillus* inhibited *Botrytis* growth; however, some of the fungal population survived after one week and even one month of interaction (**Fig. 1A**), an observation that suggests the existence of a defense mechanism activated by *Botryti*s to persist in a chemically hostile environment. Moreover, rather than simply being the attacker, the *Bacillus* population also experiences significant mortality during this interaction, particularly the bacterial cells that are in close contact with fungal hyphae (**Fig. 1B**). These findings, indicative of bidirectional antagonism, challenge the conventional view of this interaction as strictly unidirectional and point toward a more balanced relationship wherein both organisms may employ adaptive defense mechanisms. Indeed, in the long term (one week), the total *Bacillus* population was comparatively larger than that in *Bacillus* monoculture (**Fig. 1C**), despite the higher percent mortality of *Bacillus* cells attached to *Botrytis* hyphae. This result indicates that individual bacterial cells are more likely to die when they are in close contact with *Botrytis*, although the overall *Bacillus* population increases during the interaction, possibly due to a beneficial environmental niche or nutrient availability created by the presence of the fungus.

**Figure 1.**
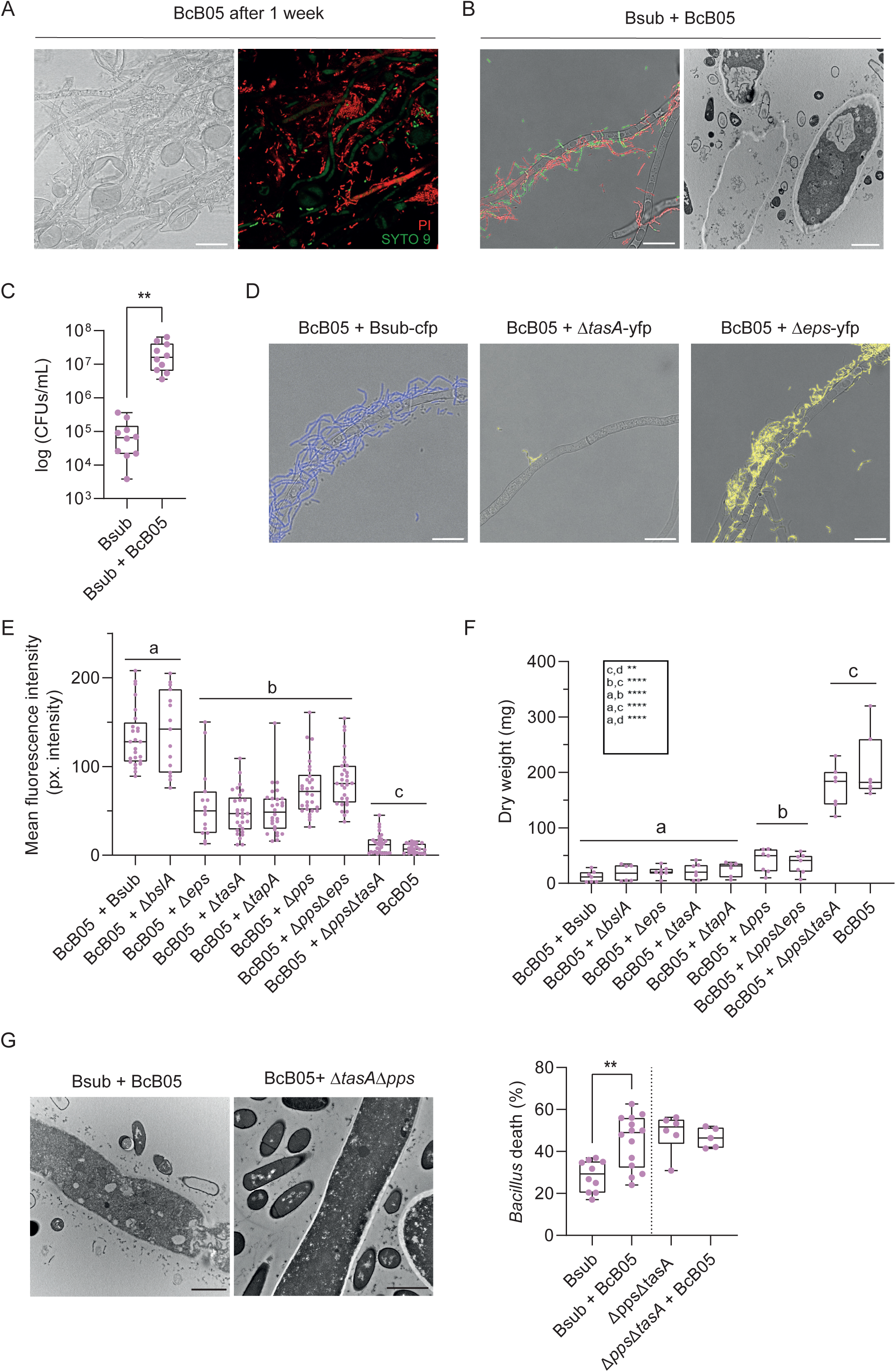
Bidirectional damage and adaptive responses in the interaction between *B. subtilis* and *B. cinerea*. **A)** Representative confocal microscopy images of *B. cinerea* after coculture with wild-type *B. subtilis* for one week. **B) Left**: confocal microscopy images showing *B. subtilis* mortality near *B. cinerea* hyphae after 24 h, visualized using live/dead staining, highlighting significant bacterial cell death near fungal structures, indicative of bidirectional aggression. **Right:** transmission electron microscopy images showing *B. cinerea* cell leakage and *B. subtilis* ghost cells surrounding fungal hyphae. **C)** Quantification of the total *B. subtilis* population in coculture after one week, demonstrating an overall increase compared with that of the monoculture. **D)** Representative images showing impaired adhesion of *B. subtilis* mutants lacking tapA or tasA to *B. cinerea* hyphae. **E)** Quantification of ROS levels in *B. cinerea* during coculture with wild-type *B. subtilis* (Bsub), mutants lacking structural and nonstructural ECM components (Δpps, ΔtasA, ΔtapA, Δeps, and ΔbslA) and double mutants (Δpps Δeps and ΔppsΔtasA). **F)** Fungal growth of *B. cinerea* during coculture ON assays. **G)** Quantification of the percentage of *B. subtilis* cell death in monoculture and during coculture with *B. cinerea*, including comparisons with the ΔtasAΔpps mutant strain, showing the influence of these ECM components on fungus-induced bacterial mortality. Scale bars for confocal microscopy represent 20 µm, and scale bars for TEM images represent 2 µm. The whisker plot shows all the measurements (pink dots), medians (black line), and minimum and maximum values (whisker ends). Statistical analyses were performed on at least three biological replicates. Statistical significance was assessed using a t test, with two asterisks indicating significant differences at P < 0.01.

To investigate the mechanism underlying the dynamic interaction between *Bacillus* and *Botrytis*, we initially examined the transcriptomic profiles of both organisms after six hours of coculture. Dual RNA-seq analyses revealed deregulation of key pathways involved in glutathione metabolism, secondary metabolite biosynthesis, or phospholipid metabolism in *Botrytis* (**Extended Data Fig. 1A**). Specifically, we observed the overexpression of *Botrytis* genes related to the fungal matrix, microbody lumen, cell wall, peroxisomes, plasma membrane, and peroxisomal matrix (**Extended Data Fig. 1B).** These results suggest the activation of a defensive strategy to preserve structural integrity and manage putative cellular damage associated with oxidative stress. Moreover, we observed notable downregulation of genes associated with the cytoskeleton, further indicating potential reorganization or damage of the internal structures of *Botrytis* due to the interaction with *Bacillus (***Extended Data Fig. 1C**). In the interaction, genes involved in secondary metabolite production (**Extended Data Fig. 2**) and genes related to *Bacillus* ECM biosynthesis (TasA, TapA, EPSs, and BslA) were upregulated in the *Bacillus* population. These results, along with the relevance of the ECM in *Bacillus* ecology and communication with other biological entities, led us to investigate the role of structural ECM components in the colonization of fungal hyphae and the antagonism toward *Botrytis*. Adhesion to hyphae was clearly impaired in mutants lacking *tapA* or *tasA* (**Fig. 1D**). Interestingly, the removal of any ECM component, including TasA, TapA or EPSs but not BslA, resulted in a reduction in reactive oxygen species (ROS) levels in *Botrytis* (**Fig. 1E**). However, no reduction in the mortality rate or fungal biomass of *Botrytis* during coculture with ΔtapA or ΔtasA strains was detected (**Fig. 1F**), likely because of the higher levels of fengycin production reported for the ΔtasA mutant^31^. The plipastatin produced by *B. subtilis* is classified in the fengycin family because of its similar structure^32^. Therefore, herein, we will refer to both as fengycin, including when discussing *Bacillus-*derived compounds. The lack of adhesion observed in the ΔtapA mutant, coupled with reduced ROS levels, was intuitively attributed to the inability of this mutant to efficiently expose TasA on the cell surface^13,33^. Compared with *Bacillus* WT cells, the Δpps mutant, unable to synthesize fengycin, triggered lower levels of ROS production in *Botrytis* (**Fig. 1E**). The ΔtasAΔpps double mutant strain lost practically all of its antifungal activity, which correlated with levels of ROS comparable to those in untreated *Botrytis* hyphae (**Fig. 1E, 1F**). Therefore, TasA and fengycin seem to synergistically influence the antagonistic activity of *Bacillus* cells toward *Botrytis*. According to our initial observation of mutual aggression, coinoculation of *Botrytis* with the ΔtasAΔpps double mutant strain completely abolished the mortality of *Bacillus* induced by *Botrytis* (**Fig. 1G**), a finding that supported a fungal offensive response activated upon perception of the ECM component TasA or fengycin.

### Lysophosphatidylcholine and ROS accumulation in *Botrytis* hyphae is correlated with the presence of TasA

Considering the relevance of the ECM in the antagonistic interaction of *Bacillus* and *Botrytis* and the differences in *Bacillus* mortality on the basis of the presence of certain ECM components, we aimed to elucidate potential metabolic changes that might modulate the interaction between these two microorganisms. We conducted a time-course comparative analysis of *Bacillus* strains (WT and derivative ECM mutants) and *Botrytis* metabolomes in monocultures and pairwise interactions at 6, 24, and 48 h. Non-targeted metabolomics analysis revealed that multiple metabolites, including glycerophosphocholines, glycerophosphoethanolamines, cyclic lipopeptides, dipeptides, tripeptides, and alkaloids, were differentially accumulated during the interaction (**Extended Data Fig. 3A**). *Bacillus* notably presented elevated levels of cyclic lipopeptides and glycerophosphoethanolamines, both of which are known for their antagonistic activity and their role in membrane stress responses^7,34,35^. In contrast, changes in glycerophosphocholine levels in *Botrytis* indicated active membrane remodeling in response to *Bacillus*, potentially aiding in adaptation to environmental stresses^36–40^. Additionally, the accumulation of dipeptides, tripeptides, and alkaloids, which are typically associated with stress and intercellular signaling, suggested a reorganization of metabolic pathways, potentially influencing the growth and survival of both organisms (**Extended Data Fig. 3A, Extended Data Fig. 3B**). Principal component analysis of the cell fraction after 24 h of interaction revealed distinct clustering patterns between *Botrytis* treated with the *Bacillus* WT or single ECM mutants. Consistent with the proposed prominent role of TasA and fengycin in the antagonistic activity, the metabolomes of *Botrytis* cocultured with the Δpps, ΔtasA, or ΔtapA mutants clustered closely with those of *Botrytis* grown in monoculture, whereas the *Bacillus* WT and mutant lacking EPSs formed a separate cluster (**Fig. 2A**). These results suggest that the interaction with *Bacillus* WT or the Δeps mutant induces substantial shifts in the *B. cinerea* metabolome. In contrast, the metabolome of *Botrytis* treated with the ΔtapA, ΔtasA, or Δpps mutant showed minimal changes and closely resembled that observed without treatment, indicating the critical contribution of TasA and fengycin in driving the metabolic modulation of *Botrytis* during the interaction.

**Figure 2.**
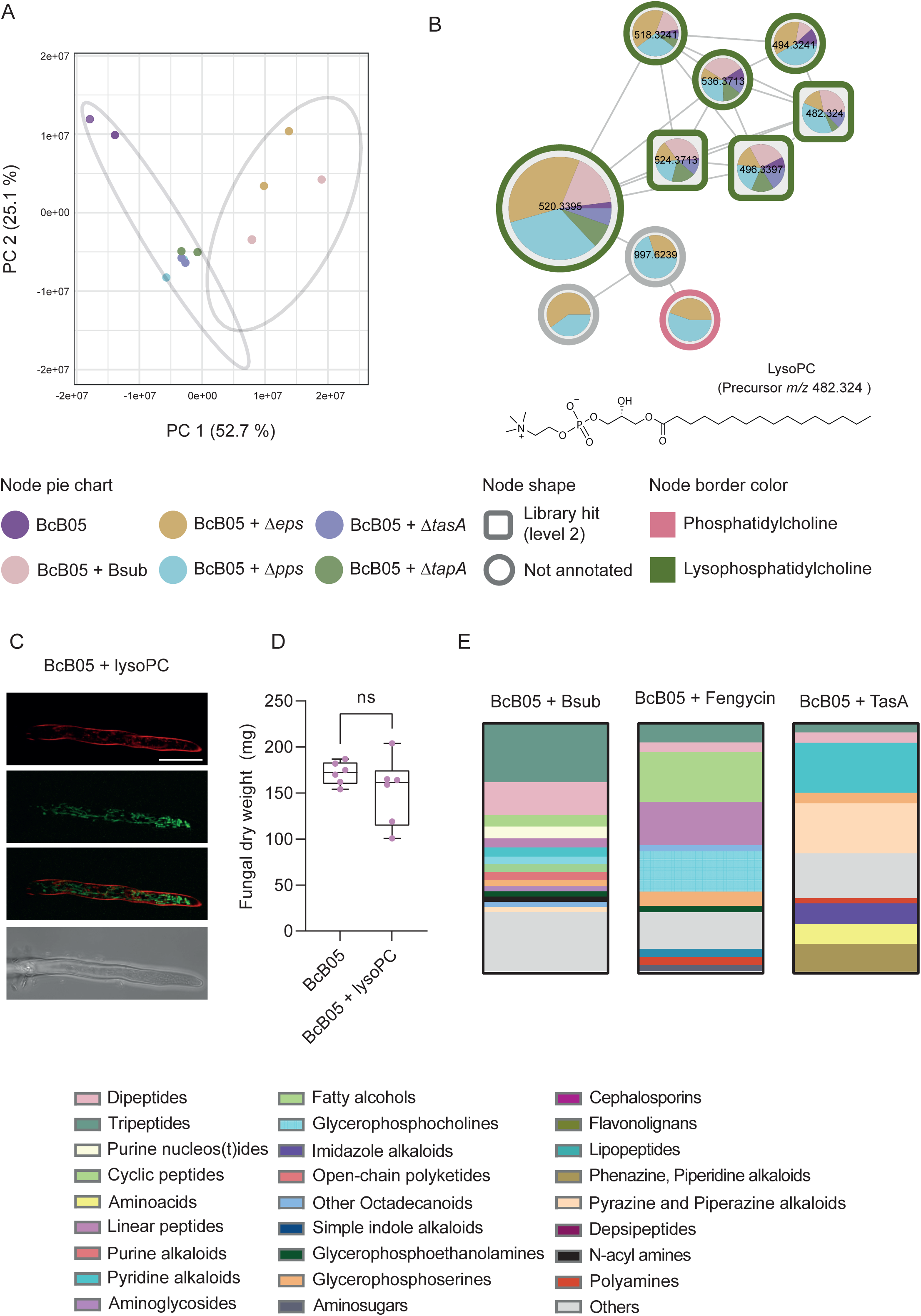
ECM structural components of *B. subtilis* drive lysophosphatidylcholine accumulation and oxidative stress in *B. cinerea*. **A)** PCA 2D score plot of the metabolome of the *B. cinerea* cell fraction after 24 h of coculture with wild-type *B. subtilis* or ECM mutants (Δpps, ΔtasA, ΔtapA, and ΔEPS). *B. cinerea* cocultured with the Δpps, ΔtasA, or ΔtapA mutant clustered closely with the *B. cinerea* monoculture control, whereas *B. cinerea* cocultured with wild-type *B. subtilis* and the Δeps mutant formed distinct clusters. The percentage of variation explained by each principal component is indicated on the axes. **B)** Molecular family of features differentially abundant in the cell fraction of *B*. *cinerea* after 48 h of treatment with wild-type *B. subtilis* or ECM mutants (Δpps, ΔtasA, ΔtapA, and ΔEPS), annotated as lysophoshphocholines according to NPClassifier. The chemical structures of annotated features and their average masses on the basis of spectral matches to GNPS libraries are also represented for the corresponding molecular families. Pie charts indicate the peak abundance of each metabolite under the corresponding conditions. The node shape indicates the level of identification according to ref.^101^. Lyso-PC accumulation is significantly reduced in interactions with ΔtasA or ΔtapA mutants, suggesting a dependence on ECM structural components. **C)** ROS levels in *B. cinerea* cultures treated with 150 µM commercial lyso-PC, confirming its role in triggering oxidative stress. **D)** Fungal growth of *B*. *cinerea* treated with 150 µM commercial lyso-PC and control *Botrytis*, showing that lyso-PC-induced ROS do not lead to *B. cinerea* growth inhibition. The whisker plot shows all the measurements (pink dots), medians (black line), and minimum and maximum values (whisker ends). For all experiments, the results of at least three biological replicates are shown. Statistical significance was assessed via a t test. **E)** Stacked bar plots represent the relative abundance of chemical classes for the top 250 features significantly increased in the *B. cinerea* cell fraction after 24 h of treatment with TasA, fengycin, or wild-type *B. subtilis*, compared with untreated *B. cinerea*. The features were ranked using volcano plots generated in MetaboAnalyst. Metabolite chemical class prediction was performed using SIRIUS, and the metabolites were classified with NPClassifier. Scale bars represent 20 µm.

Further analysis of the metabolomes revealed notable accumulation of lysophosphatidylcholine (lyso-PC) in the *Botrytis* cell fraction after coculture with *Bacillus*. Lyso-PC is a lipid signaling molecule associated with ROS generation, cellular damage and inflammatory processes in other biological systems^40–45^. Interestingly, lyso-PC was practically absent in *Botrytis* treated with the ΔtasA or ΔtapA mutant, a finding that led us to correlate the accumulation of lyso-PC with the presence of *Bacillus* ECM components (**Fig. 2B**). We hypothesize that the accumulation of lyso-PC might correlate with the levels of ROS accumulated in *Botrytis* during the interaction (**Fig. 1E**). Commercial lyso-PC exogenously added to *Botrytis* cultures triggered significant increases in ROS levels (**Fig. 2C, Extended Data Fig. 3C**), confirming that lyso-PC contributes to the oxidative stress observed during the *Bacillus-Botrytis* interaction, a physiological response triggered by the presence of the *Bacillus* ECM structural component TasA. However, the level of ROS produced after the addition of lyso-PC did not appear to lead to fungal death or growth inhibition (**Fig 2D**), suggesting that lyso-PC may function primarily in signaling rather than inhibiting the growth of *Botrytis*. According to this finding, the external addition of purified TasA to *Botrytis* reproduced the accumulation of ROS observed in previous coculture experiments and promoted the accumulation of lyso-PC in *Botrytis* cells. Fengycin also triggered ROS accumulation in *Botrytis* hyphae; however, the level of lyso-PC accumulation did not increase significantly.

To better understand the specific metabolic changes triggered by the most active ECM components, we analyzed the metabolome of *B. cinerea* treated with purified TasA or fengycin. Chemical class analysis of non-targeted metabolomics data with CANOPUS^46^ revealed that during the interaction of *Botrytis* with *Bacillus*, glycerophosphocholines, dipeptides, tripeptides, cyclic peptides, and alkaloids accumulated after 24 h, as shown in **Extended Data Fig. 3B**. Fengycin-treated *Botrytis* exhibited accumulation of dipeptides, tripeptides and glycerophosphocholines, which is consistent with its membrane-disrupting activity. Similarly, TasA-treated *Botrytis* presented minor accumulation of dipeptides and tripeptides compared with fengycin-treated *Botrytis*, as well as pronounced accumulation of alkaloids, particularly piperazine and pyridine alkaloids **(Fig. 2D**). These findings suggest that dipeptides and tripeptides accumulate in the presence of both ECM components. Glycerophosphocholine production is driven by fengycin, and alkaloid production is associated with TasA. This differential response of *Botrytis* at the metabolic level led us to investigate specific physiological and anatomical damage induced by these molecules produced by *Bacillus*.

### TasA modifies the structural integrity of *Botrytis* hyphae, compromising virulence

*Botrytis* accumulates ROS in response to the presence of *Bacillus* ECM; however, evident inhibition of fungal growth or accumulation of lyso-PC was observed in the presence of fengycin or TasA, respectively. The addition of purified TasA to *Botrytis* cultures promoted a curling morphology of hyphae (**Fig. 3A**), a macroscopic phenotype reasonably associated with disorganization of the fungal cell surface. To examine the hypothetical impact of TasA on the biophysical properties of the fungal matrix, we conducted magnetic resonance imaging (MRI) analyses to assess T2 relaxation times across different colony regions, which reflect water content and diffusion. High-resolution images of untreated *Botrytis* macrocolonies revealed significant heterogeneity, with varying water contents and relatively high T2 relaxation times in the center of the colonies. In contrast, *Botrytis* treated with TasA presented a homogeneous distribution, with uniformly lower T2 times. This finding suggested the existence of an intact matrix in *Botrytis* colonies that is able to retain water within the colony and that disruption of matrix integrity in the presence of TasA reduces water retention and thus structural organization (**Fig. 3B**).

**Figure 3.**
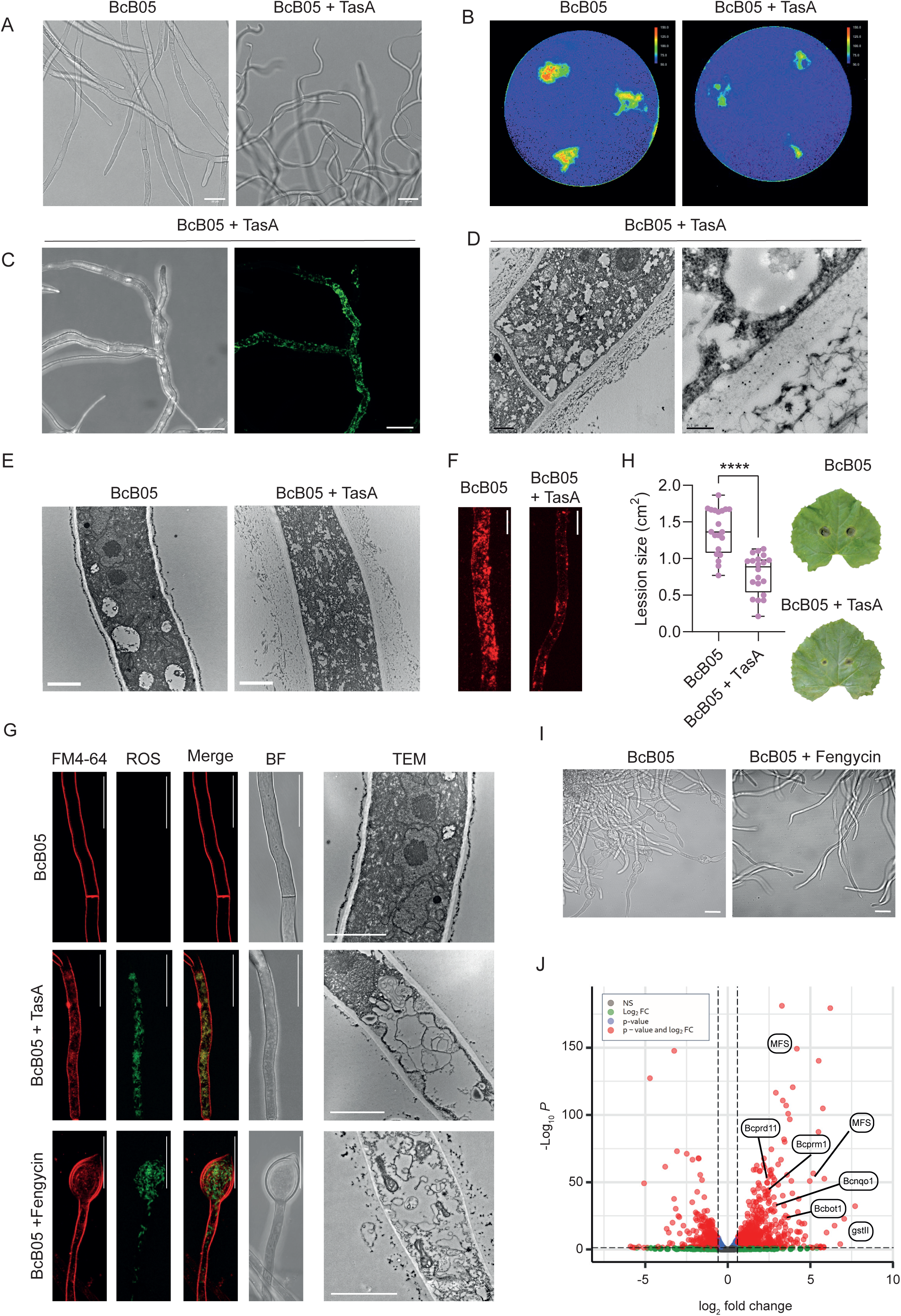
TasA and fengycin differentially impact *B. cinerea* physiology. **A)** Macroscopic phenotypic analysis showing the curling morphology of *B. cinerea* hyphae upon treatment with purified TasA, likely due to disorganization of the fungal cell surface. Scale bars represent 20 µm. **B)** Magnetic resonance imaging-based T2 relaxation time analysis showing that untreated *B. cinerea* macrocolonies retain water heterogeneously, with higher T2 times at the colony center. In contrast, TasA-treated colonies presented a uniform distribution of water content and relatively low T2 relaxation times, suggesting disruption of fungal matrix integrity and reduced water retention. **C)** Immunocytochemistry using anti-TasA antibodies demonstrating extensive decoration of fungal hyphae after *B. cinerea* treatment with 3 µM purified TasA. Scale bars represent 20 µm. **D)** Transmission electron micrograph of negatively stained thin sections of *Botrytis* hyphae with immunogold labeling revealing TasA accumulation in the outer and inner layers of the fungal cell wall, which is primarily composed of chitin and chitosan. The scale bar represents 1 μm for the image on the left and 0.2 μm for the zoomed-in image on the right. **E)** TEM images of untreated hyphae display a sharply defined β-glucan layer outside the cell wall (left), whereas TasA-treated hyphae exhibit disorganized β-glucan structures (right). The scale bar represents 2 μm. **F)** Tubulin staining of *B. cinerea* hyphae revealed significantly fewer tubulin foci in TasA-treated samples than in control samples, indicating cytoskeletal disorganization. **G)** Confocal microscopy images of ROS levels in 3 μM TasA-treated *B. cinerea* hyphae, obtained using double staining with FM4-64 to stain the membrane, and TEM images showing ultrastructural damage, including autophagosome formation, in TasA-treated *Botrytis*. Scale bars: 20 μm for confocal microscopy images and 2 μm for TEM images. **H)** *In planta* experiments showing reduced lesion size and fungal colonization in plants treated with TasA compared with those in untreated controls. The whisker plot shows all the measurements (pink dots), medians (black line), and minimum and maximum values (whisker ends). For all experiments, the results from at least three biological replicates are shown. Statistical significance was assessed via a t test, with quadruple asterisks indicating significant differences at P < 0.0001. **I)** Brightfield microscopy of *B. cinerea* hyphae demonstrating extensive chlamydospore formation upon fengycin treatment (lower panel) compared with the control treated with methanol (upper panel). Scale bars equal 20 µm. **J)** Volcano plot of DEGs identified by RNA-seq in *B. cinerea* treated with fengycin for 6 hours and untreated hyphae. P values were calculated on the basis of the Fisher method using nominal P values provided by edgeR and DEseq2. The dashed lines represent the thresholds defined for P (horizontal) and the fold change (vertical) for a gene to be considered a DEG. Genes related to virulence and detoxification pathways are labelled as follows: MFS (major facilitator superfamily), gstII (glutathione S-transferase), Bcprd11 (peroxidase, involved in the response to oxidative stress), Bcprm1 (serine peptidase, involved in fungal survival), Bcnqo1 (oxidoreductase) and Bcbot1 (botrydial biosynthesis gene, involved in secondary metabolism).

The relevance of TasA for the cellular attachment of *Bacillus* cells to *Botrytis* hyphae led us to initially evaluate the role of this protein in the disorganization of the fungal cell wall. Immunocytochemistry analysis using anti-TasA antibodies revealed that the fungal hyphae were extensively decorated with the fluorescent signal associated with TasA (**Fig. 3C**). Transmission electron microscopy (TEM) analysis of thin sections of *Botrytis* hyphae and immunogold labeling confirmed the accumulation of TasA-related signals in the outer and inner fungal cell wall layers, which contained mainly chitin and chitosan, the two major components of the fungal cell wall (**Fig. 3D**). TEM control images of *B. cinerea* are shown in **Extended Data Fig. 4A**. The affinity of TasA for both chitin and chitosan was biochemically demonstrated via polysaccharide affinity assays (**Extended Data Fig. 4B**). In the TEM analysis, however, any alteration in the chitin-chitosan layer was disregarded, and noticeable disorganization of the β-glucan layer of the fungal cell wall was observed following treatment with TasA, which, in untreated hyphae, appeared as a sharply defined electrodense line outside the less electrodense chitin-chitosan layer (**Fig. 3E**).

Fungal shape and structural integrity are largely regulated by the cytoskeletal element actin filaments and microtubules^47^, and disruption of the cytoskeleton may induce a curling morphology. Therefore, we reasoned that disorganization of the β-glucan layer induced by TasA might cause cytoskeleton dysfunction, weakening the structural integrity of the fungal cell wall and compromising the ability of *Botrytis* to maintain its typical hyphal architecture and function. Diverse lines of experimental evidence support this hypothesis: i) downregulation of genes related to cytoskeletal organization in transcriptomic analysis of TasA-treated *Botrytis* hyphae (**Extended Data Fig. 4C**), ii) a significant decrease in tubulin foci of hyphae of *Botrytis* treated with TasA upon specific staining of the cytoskeleton with a tubulin tracker (**Fig. 3F, Extended Data Fig. 4D**), and iii) cytoplasmic damage characterized by autophagosome formation observed via TEM analysis of thin sections of *Botrytis* hyphae (**Fig. 3G**). Collectively, these findings demonstrate that in addition to facilitating the physical contact of *Bacillus* cells with *Botrytis* hyphae, TasA disrupts the physical and molecular architecture of *Botrytis* cells. These results confirm the active participation of TasA in the arsenal of molecules secreted by *Bacillus* to efficiently antagonize *Botrytis.* The failure of EPSs to induce significant changes at the ultrastructural level (**Extended Data Fig. 5A**) is likely related to the limited access of this bacterial polymer to the fungal cell wall or membrane. However, the increase in ROS levels suggested that *Botrytis* can sense the presence of EPS, most likely triggering a differential response to TasA (**Extended Data Fig. 5B**). We further demonstrated in an *in vivo* plant model (melon) that the structural disorganization and morphological changes triggered by TasA translated into a reduced ability of *Botrytis* to induce symptoms in leaves (**Fig. 3H**).

Fengycin is known to target fungal membranes, and consistent with this, the cytoplasmic disorganization of *Botrytis* hyphae was more pronounced than that induced by TasA. The most noticeable cytological alterations were autophagosome formation (**Fig. 3G**), plasmolysis, and extensive formation of chlamydospores (**Fig. 3I**), which are differentiated fungal cells associated with resistance to external aggressions^48–50^. Consistent with these cellular reorganizations, transcriptomic analysis of *Botrytis* treated with fengycin after 6 h revealed upregulation of virulence-related genes and detoxification pathways, particularly those associated with glutathione, oxidoreductases, and major facilitator superfamily (MFS) transporters, indicating the activation of a chemical defense mechanism oriented toward mitigating the oxidative stress and cytological damage inflicted by fengycin (**Fig. 3J**). All the ROS quantifications for *B. cinerea* with purified ECM components are provided in **Extended Data Fig. 4D**.

### Antibacterial oxylipins are differentially produced by *Botrytis* in response to TasA or fengycin

We determined how *Botrytis* hyphae (cell fraction, **Extended Data Fig. 2E**) respond differentially metabolically and anatomically to TasA or fengycin, two active *Bacillus* ECM components involved in the interaction. However, the factors delivered by *B. cinerea* that are lethality to *B. subtilis* cells (**Fig. 1B**) remain elusive. To illuminate the change in metabolite production associated with hyphal damage in *B. cinerea*, we performed non-targeted metabolomics analyses of the *Botrytis* supernatant at different time points (6, 24, and 48 h) after treatment with purified TasA or commercial fengycin. Consistent with the morphological changes (**Fig. 3G**), our data revealed that TasA and fengycin differentially impacted *Botrytis* physiology, the metabolome, and most likely the fungal lifestyle. Fengycin treatment led to substantial secretion of metabolites by *Botrytis* into the media, particularly after 24 h, and TasA-treated *Botrytis* reduced the overall accumulation of metabolites (**Extended Data Fig. 6**). The pool of accumulated metabolites in the fungal supernatant fraction under both treatments included oxygenated fatty acids or oxylipins, especially after treatment with fengycin **(Fig. 4A**). Oxylipins are derived from linoleic and linolenic acids and are known as developmental regulators or mediators of interspecies communication across various biological systems, including mammals, bacteria, fungi, and plants^51–53^. In filamentous fungi, oxylipins constitute an essential defensive mechanism for fungal adaptation to the environment^54–56^. The linolenic-derived oxylipins detected included stearidonic acid (SDA), 9-OxoOTrE, trans-EKODE, 13-OxoODE and 9S-HOTrE, and the linoleic-derived epoxides detected included 12,13-EpOME and 9(10)-EpOME (**Extended Data Fig. 7**). One of the most abundant oxylipins, SDA, exhibited potent antibacterial effects, significantly contributing to *B. subtilis* mortality at concentrations as low as 6 µg/mL. This finding endorses SDA as a potential key factor in the defensive arsenal of *Botrytis* activated upon the differential cellular aggression inflicted by the *Bacillus* ECM molecule fengycin and, to a lesser extent, TasA.

**Figure 4.**
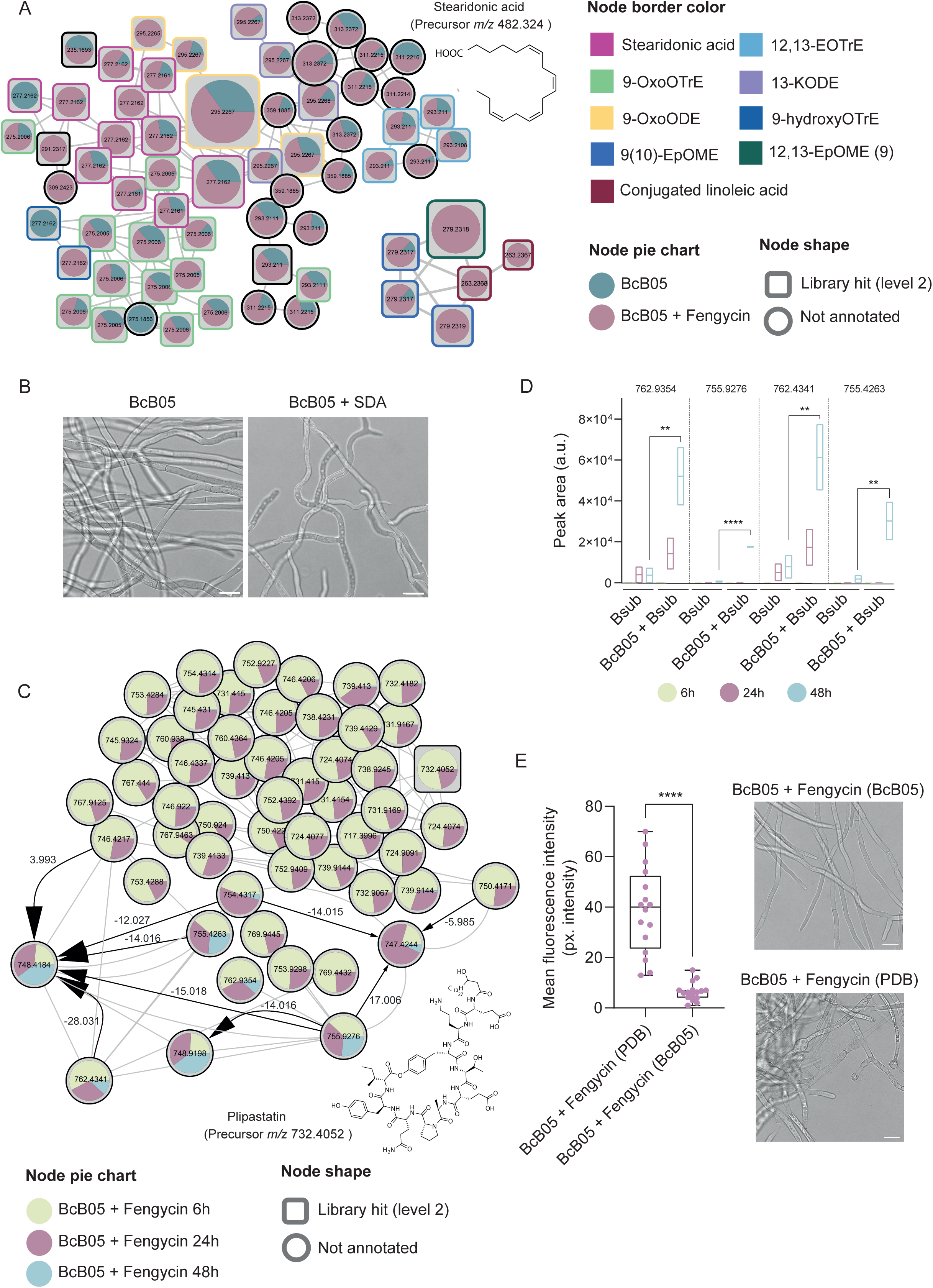
Counteracting *B. subtilis* aggression: oxylipin production and fengycin enzymatic degradation. **A)** Molecular families of features differentially abundant in the supernatant fraction of *B. cinerea* after 24 h of treatment with 10 μM fengycin or no treatment. The chemical structures of annotated features and their average masses on the basis of spectral matches to GNPS libraries are also represented for the corresponding molecular families. Pie charts indicate the peak abundance of each metabolite under the corresponding conditions. The node shape indicates the level of identification according to ref.^101^. The node border color indicates the compound name according to the GNPS library. **B)** Representative confocal microscopy images of untreated *B. cinerea* (left) or *B. cinerea* treated with 1 µg/mL commercial SDA (right), triggering hyperbranching. Scale bars equal 10 µm. **C)** Molecular family analysis of fengycin and its structural variants in *B. cinerea* supernatants after the addition of 10 µM fengycin, which were sampled at 6, 24, and 48 h. Pie charts represent the mean peak abundance of metabolites at each time point, with node shapes indicating the level of metabolite identification. The arrows indicate the chemical directionality of the modifications observed, with arrow sizes corresponding to ChemProp scores. The labels above the arrows reflect the mass differences between related metabolites. **D)** Floating bar plots showing the peak abundances of four selected putative degradation products of fengycin in *B. subtilis* cultures growing alone and in coculture with *B. cinerea* at 6, 24, and 48 h. **E) Left:** Quantification of ROS levels in *B. cinerea* after treatment with fengycin preincubated in PDB medium for 24 h versus fengycin preincubated with *Botrytis* for the same amount of time. Lower ROS levels were detected in *B. cinerea* treated with fengycin preincubated with *B. cinerea*, suggesting neutralization of fengycin activity. The whisker plot shows all the measurements (pink dots), medians (black line), and minimum and maximum values (whisker ends). For all experiments, the results from at least three biological replicates are shown. Statistical significance was assessed via a t test, with quadruple asterisks indicating significant differences at P < 0.0001. **Right:** Confocal microscopy images showing cytological damage and chlamydospore formation in *B. cinerea* treated with fengycin preincubated with PDB as a control, whereas *B. cinerea* treated with fengycin preincubated with *B. cinerea* presented no apparent cytological damage. Scale bars equal 20 µm.

In addition to their antibacterial activity, fungus-derived oxylipins seem to mediate intrapopulation communication within fungi. Accordingly, SDA exogenously added to *Botrytis* culture induced hyphal hyperbranching (**Fig. 4B**). Hyperbranching in filamentous fungi is perceived as an ecological strategy that enhances the ability of fungi to explore and capture nutrients, as well as provides protection against threats. This behavior is part of an interference competition strategy that increases the likelihood of survival and fitness in resource-limited environments^57^. Therefore, the production of these oxylipins by *Botrytis* seems to exemplify a dual function: damaging competing microbes such as *Bacillus* while promoting alternative adaptive growth strategies.

### *Botrytis* enzymatically degrades fengycin, neutralizing its antifungal activity

Two alternative strategies could explain the long-term resistance of *Botrytis* to the physicochemical effects of *Bacillus*: i) antagonism toward *Bacillus* cells (**Fig. 1B)** and ii) neutralization of the toxicity of the antifungal molecules produced by *Bacillus*. We have proposed oxylipins as one type of antibacterial agents secreted by *Botrytis* in response to the action of fengycin, with a deleterious effect on *Bacillus* cells (**Fig. 4A**); thus, we focused on demonstrating the implications of the second strategy. Considering the relevance of fengycin in the antifungal activity of *B. subtilis*, we analyzed the degradation of this lipopeptide during the interaction. By leveraging non-targeted metabolomics data from serial incubation with fengycin, we analyzed proportionality changes (anticorrelation behavior) in chemically related metabolites through molecular networking and the ChemProp2 computational tool^58^. This analysis revealed a significant reduction in fengycin levels in the cell-free supernatant 24 to 48 h after the addition of commercial fengycin to *Botrytis* cultures. This reduction in the original fengycin level was accompanied by the accumulation of specific structural variants of fengycin with *m/z* values of 762.9354, 755.9276, 762.4341 and 755.4263, likely representing degradation products (**Fig. 4C**). Similar dynamics were observed for native fengycin produced by *Bacillus* upon interaction with *Botrytis*, indicating the ability of *Botrytis* to chemically modify fengycin (**Fig. 4D**). To evaluate the biological implications of this degradation, *Botrytis* was incubated with commercial fengycin for 24 h, after which the supernatant was added to fresh *Botrytis* cells. Fengycin exposed to *Botrytis* lost its antifungal activity, as demonstrated by the absence of fungal growth inhibition, lack of cellular damage, and lack of increase in ROS levels, compared with that of control fengycin incubated in PDB simultaneously (**Fig. 4E**). Two complementary observations supported the active enzymatic degradation of fengycin carried out by *Botrytis* during the interaction: i) the fact that the pH of the culture did not change significantly and ii) the specific fengycin degradation products with known enzymatic degradation profiles reported in the literature^59^. To identify putative enzymes involved in the chemical transformation of fengycin, we analyzed the genes that were upregulated in *Botrytis* following fengycin treatment or upon interaction with wild-type *Bacillus* via a previously described workflow involving RNA-Seq data^60^. We focused on proteins predicted to be extracellular and conducted a search for conserved protein domains to uncover key functional motifs involved in their activity, identifying putative enzymes involved in fengycin degradation (**Suppl. Table 1**). This is a notable observation since the degradation of lipopeptides is reported to occur during competition with other bacteria^61,62^ but not fungi. This finding indicates that there could be an additional defensive mechanism used by *Botrytis* and probably other filamentous fungi to neutralize antifungal molecules produced by *Bacillus*.

### Degradation of fengycin coincides with the secretion of fungistatic bacillaene by *Bacillus*

Following the degradation of *Bacillus* primary antifungal compounds with activity against *Botrytis*, metabolomic analyses revealed an increase in the production and accumulation of bacillaene, another secondary metabolite from the *Bacillus* chemical arsenal^63,64^. Specifically, bacillaene accumulated at significantly greater levels in the supernatants of *Bacillus* cells cocultured with *Botrytis* than in those of *Bacillus* cells grown in monoculture after 24 h (**Fig. 5A**). Non-targeted metabolomic analysis of the interaction of *Botrytis* with the Δpps mutant (unable to produce fengycin) revealed elevated accumulation of bacillaene compared with the levels accumulated during the interaction with wild-type *Bacillus*, indicating compensatory regulation between fengycin and bacillaene (**Extended Data Fig. 8**). This finding indicates a flexible defensive strategy adopted by *Bacillus* that involves the use of bacillaene in response to the fengycin-neutralizing activity of *Botrytis*.

**Figure 5.**
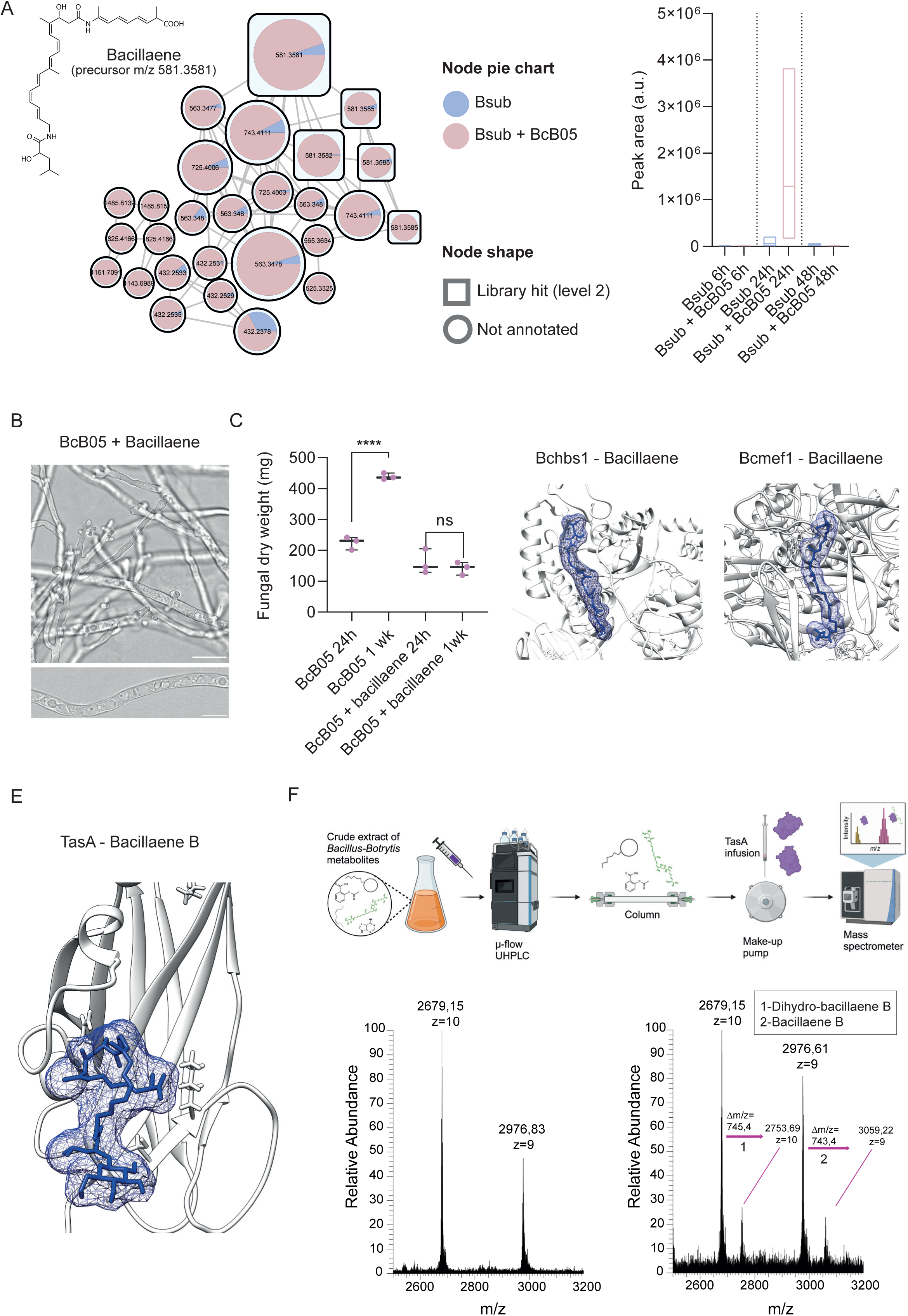
Interaction with *B. cinerea* triggers bacillaene production, revealing its fungistatic effects. **A) Left:** Molecular family analysis of bacillaene in *B. cinerea* supernatant after interaction with *B. subtilis* compared with that in the *B. subtilis* monoculture at 24 h. Pie charts represent the mean peak abundance of metabolites in the supernatant, with node shapes indicating the level of metabolite identification according to GNPS libraries. **Right:** Floating bar plots showing the peak abundance of bacillaene (precursor m/z 581.3581) in *B. subtilis* supernatants from monoculture and coculture with *B. cinerea* at 6, 24, and 48 h, highlighting bacillaene accumulation during interactions at 24 h. **B)** Confocal microscopy images of *B. cinerea* treated with 60 µg/mL purified bacillaene for 24 h, showing cytoplasmic vacuolization and hyphal tip splitting. Scale bars represent 20 µm. **C)** Quantification of the fungal mass after 24 h and one week demonstrated halted fungal growth upon bacillaene treatment, confirming its fungistatic activity. The whisker plot shows all the measurements (pink dots), medians (black line), and minimum and maximum values (whisker ends). For all experiments, the results from at least three biological replicates are shown. Statistical significance was assessed via a t test, with quadruple asterisks indicating significant differences at P < 0.0001. **D)** Molecular docking analysis predicted a putative binding site between bacillaene and the cytoplasmic elongation factor Bchbs1, as well as between bacillaene and the mitochondrial elongation factor Bcmef1, in *Botrytis cinerea*. **E)** Molecular docking analysis predicted a strong binding interaction between bacillaene B (precursor m/z 743.4) and TasA. **F) Top:** Overview of the native metabolomics workflow. **Bottom:** Analyses of native metabolomics assays between purified TasA and the crude extract of metabolites. **Left:** Mass spectrum of TasA alone. **Right:** Mass spectrum of TasA in the presence of the crude metabolite extract from the *Bacillus-Botrytis* coculture, revealing a distinct mass shift corresponding to the formation of the TasA-bacillaene complex when bacillaene passed through the column. This shift indicates the binding of bacillaene A, an isoform of bacillaene functionalized with a hexose group.

To test whether this highly unstable metabolite^65,66^ affects fungal growth, *Botrytis* cultures were treated with purified bacillaene at a concentration of 60 µg/mL. Hyphal tip splitting and massive cytoplasmic vacuolization were the most notable fungal morphological changes (**Fig. 5B**). Furthermore, the external addition of purified bacillaene halted *Botrytis* growth, whereas the untreated *Botrytis* continued to grow, confirming the fungistatic role of bacillaene (**Fig. 5C**). Bacillaene has been reported to be bacteriostatic and able to disrupt bacterial protein synthesis by targeting the prokaryotic elongation factor FusA^28,67–71^. To explore whether a similar mechanism might occur in fungi, we conducted a BLAST search between *FusA* and the genome of *Botrytis*, which revealed 50% and 30% homology between the bacterial elongation factor and two *B. cinerea* elongation factors: the mitochondrial elongation factor encoded by *Bcmef1* and the cytoplasmic elongation factor encoded by *Bchbs1*, respectively. Docking studies predicted a potential ligand□protein binding interaction between bacillaene and both elongation factors, suggesting their possible role as targets of bacillaene **(Fig. 5D**). Additionally, in a previous study, we reported that *B. cinerea* treated with bacillaene developed mutations in the elongation factor gene^71^. To investigate this further, *B. cinerea* cultures were treated with bacillaene for both short (24 h) and long (1 week) durations. Whole-genome sequencing of *B. cinerea* subjected to long-term treatment revealed several mutations likely impacting the cytoplasmic elongation factor *Bchbs1*, as well as mutations in adjacent genes within the same transcript (**Suppl. Table 2**). These results suggest that *Bchbs1* is a likely target of bacillaene and demonstrate the capacity of *B. cinerea* to adapt to its presence, potentially as part of an adaptive response to mitigate the effects of bacillaene and maintain its viability.

Bacillaene is a highly unstable molecule that is particularly sensitive to light and oxygen^65,66^, which raises the question of the mechanism that preserves the functionality of this ecologically significant molecule. *Bacillus* ECM is known to retain secondary metabolites^72^, and we have shown that ECM components are upregulated during the interaction with *Botrytis*, two observations that led us to propose an ECM with a bacillaene-stabilizing role. TasA is capable of disorganizing the β-glycan layer and reaching the chitin-chitosan layer of the fungal cell wall, which further supports the hypothesis that TasA is a molecular vehicle for the stabilization and efficient delivery of bacillaene to the *Botrytis* hyphal surface. Molecular docking studies using the resolved crystal structure of TasA and bacillaene predicted the highly likely interaction of bacillaene with TasA (**Fig. 5E**). We performed native metabolomics experiments^73^ to explore the interaction between TasA and the crude extract containing metabolites from *Bacillus-Botrytis* cocultures. Native metabolomics, performed by combining non-targeted metabolomics and native mass spectrometry, showed that the fold and function of TasA were preserved^73^. The crude metabolite extract was first separated via microflow UHPLC, and subsequently (postcolumn), a make-up buffer neutralized the mobile phase, and purified TasA was infused via a T-splitter before mass spectrometry (MS) analysis (**Fig. 5F, top**). Upon bacillaene passage through the system, the interaction with TasA was monitored via MS, which revealed a molecular mass shift corresponding to the formation of the TasA-bacillaene complex (**Fig. 5F, bottom**). This molecular mass shift, along with a distinct retention time, indicated the binding of bacillaene A, a version of bacillaene functionalized with a sugar residue in its structure, to TasA. We hypothesize that binding to TasA protects bacillaene A from degradation while ensuring the stability of this metabolite and its effective delivery to the fungal target. These findings revealed an interconnection among the ECM and secondary metabolites, more specifically, TasA, as a carrier and stabilizer of structurally unstable bioactive molecules during microbial interactions.

### *B. subtilis* feeds on *B. cinerea* chitosan to maintain fengycin production

The switch from fungicidal activity mediated by fengycin to fungistatic activity driven by bacillaene suggests a shift in the strategy adopted by *Bacillus* during its interaction with *Botrytis*. This change likely leads to long-term nutritional benefits, probably in oligotrophic environments, where maintaining a balance between microbial competition and coexistence could be advantageous for *B. subtilis*. In this context, the *B. subtilis* population density increased in coculture, mostly as vegetative cells and not spores, compared with monoculture (**Fig. 1C**). One explanation for these population dynamics is associated with the ability of *B. subtilis* cells to feed on *B. cinerea* hyphal structural components as nutrients. We hypothesized that chitosan is the preferred carbon source used by *B. subtilis*, which is supported by two complementary observations: first, the accessibility of *B. subtilis* cells to chitosan is mediated by the chemical affinity of TasA (**Fig. 3D and Extended Data Fig. 4A**), and second, the existence of a *B. subtilis* chitosanase encoded by the gene *csn*, an enzyme that hydrolyses the β-1,4-glycosidic bonds of chitosan molecules^74–76^. The dynamic growth of bacteria in minimal MSGG medium with chitosan as the sole carbon source revealed the ability of the *B. subtilis* WT strain to grow; however, a Δcsn mutant strain lacking functional chitosanase was not able to grow (**Fig. 6A**). Further *in situ* degradation of fungal chitosan by *Bacillus* during the interaction with *Botrytis* was confirmed via the use of the chitosan dye eosin Y^77^ and fluorescence microscopy (**Fig. 6B**). Furthermore, the addition of purified chitosanase to *B. cinerea* culture produced a similar reduction in the fluorescence signal of fungal hyphae, indicative of chitosan loss, underscoring its role in fungal cell wall breakdown (**Fig. 6B).** Metabolomic analysis further highlighted the role of chitosanase in the interaction between *B. subtilis* and *B. cinerea*. After 24 h of coculture, we observed lower accumulation of fengycin in the supernatant of cocultures with the Δ*csn* mutant than in that of cocultures with wild-type *B. subtilis* (**Fig. 6C**). This result suggests a link between chitosanase activity, or the degradation of chitosan, and fengycin production. Flow cytometry analysis further supported this connection, revealing heightened fengycin promoter activity in the presence of chitosan after one week, which indicates that *B. cinerea* cell wall components sustain expression over time (**Fig. 6D**) and explains why, in the interaction, without accessible chitosan, fengycin accumulation is reduced. This result also suggested the possibility of a synergistic effect between Csn and fengycin against *Botrytis*. Compared with single treatments, the application of purified Csn and fengycin at the same time to *B. cinerea* significantly increased ROS production (**Fig. 6E**). These findings suggest that *B. subtilis* employs a dual strategy, nutrient acquisition and metabolic modulation, through both physical and chemical means to keep *B. cinerea* in a slowed growth state, ensuring the sustained availability of a carbon source in long-term balanced antagonistic coexistence.

**Figure 6.**
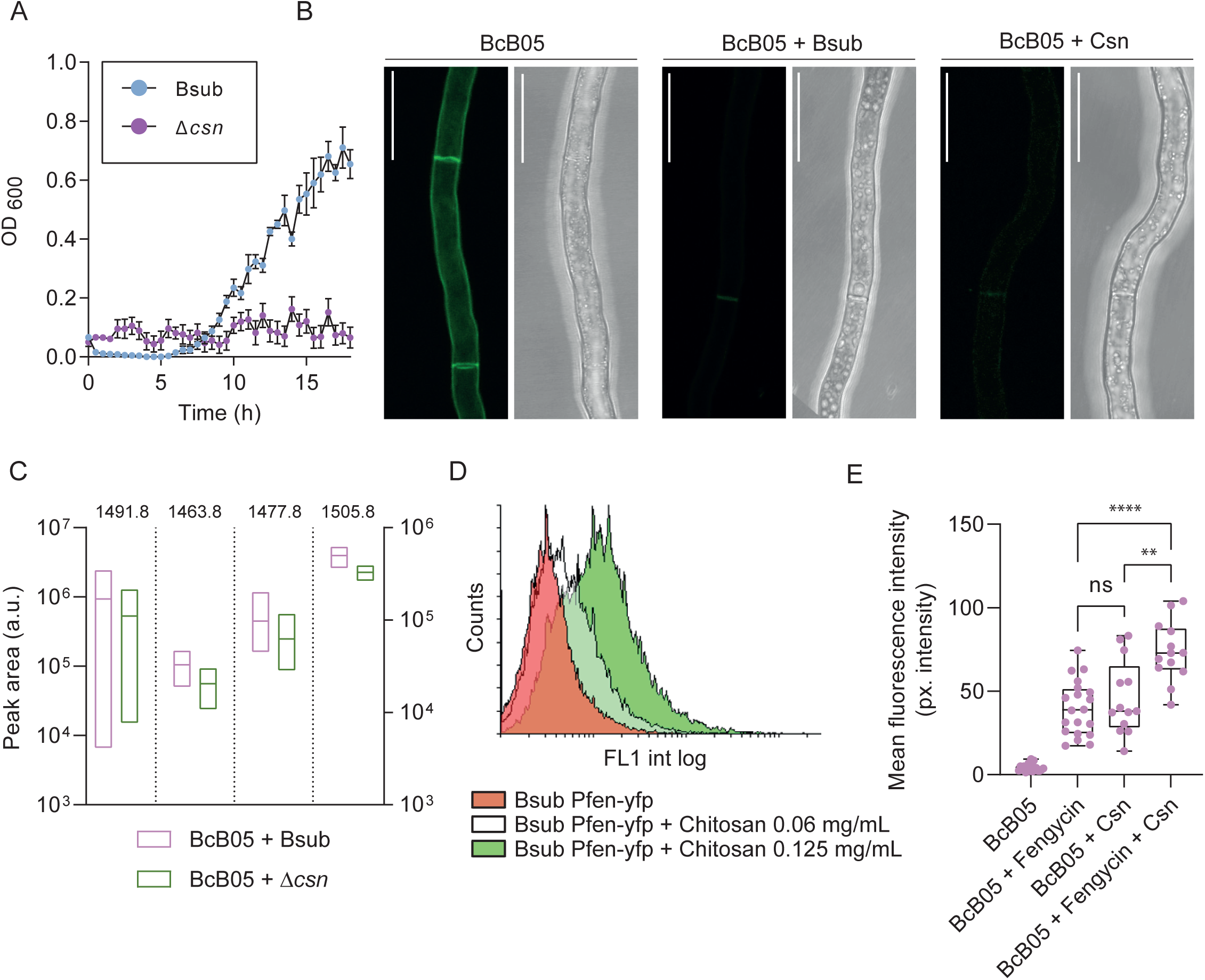
Chitosan degradation and the synergistic effects of chitosanase and fengycin contribute to *B. subtilis* growth. **A)** Growth curve of the wild-type and Δcsn *B. subtilis* strains in minimal MSGG medium supplemented with chitosan as the sole carbon source, showing no growth of the Δcsn mutant, highlighting the role of Csn in enabling *B. subtilis* to utilize chitosan. **B)** Confocal microscopy images of *B. cinerea* hyphae stained with eosin Y to visualize chitosan. The fluorescence signal was absent in hyphae treated with wild-type *B. subtilis* or 0.2 mg/mL purified Csn, demonstrating chitosan degradation. The scale bar equals 20 µm. **C)** Floating bar plot showing the peak abundances of four fengycin structural variants in the supernatant of *B. cinerea* after 24 h of interaction with wild-type *B. subtilis* or interaction with the Δcsn mutant. **D)** Flow cytometry analysis of *B. subtilis* expressing YFP under the fengycin promoter (Bsub Pfen-yfp) in the presence of control conditions, 0.06 mg/mL chitosan, or 0.125 mg/mL chitosan, demonstrating chitosan-induced maintenance of the fengycin promoter over time. **E)** Quantification of ROS intensity in *B. cinerea* treated with 10 µM fengycin, 0.2 mg/mL purified chitosanase, or a combination of both, demonstrating a synergistic effect between chitosanase and fengycin in inducing ROS accumulation. The whisker plot shows all the measurements (pink dots), medians (black line), and minimum and maximum values (whisker ends). In all experiments, at least three biological replicates are shown. Statistical significance was assessed via a t test, with double and quadruple asterisks indicating significant differences at P < 0.01 and P < 0.0001.

**Figure 7.**
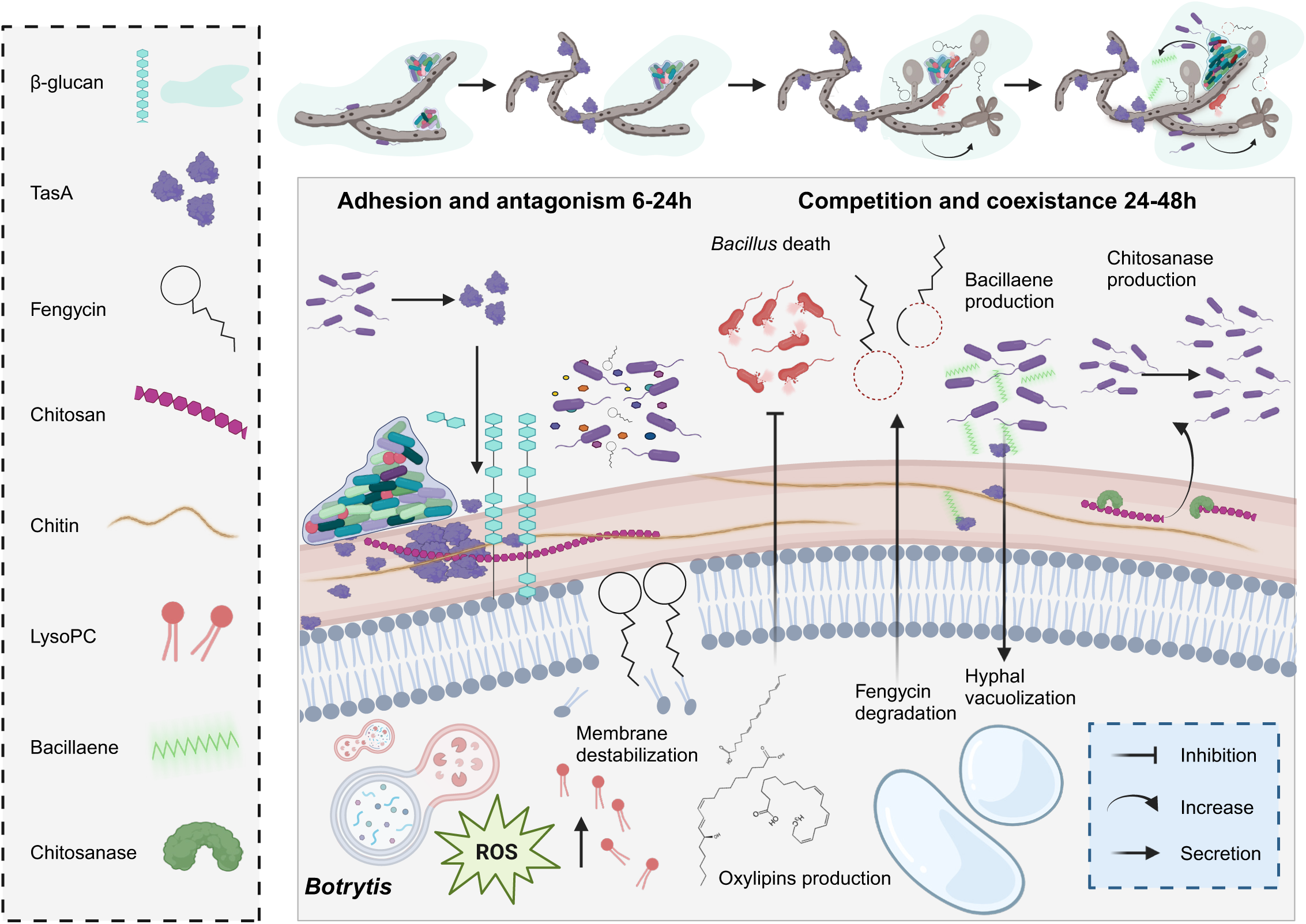
Overview of the dynamic interaction and counterdefense between *B. subtilis* and *B. cinerea*. The interaction between *Bacillus* and *Botrytis* is a highly dynamic process characterized by mutual adaptation. *Bacillus* ECM components, specifically TasA and fengycin, induce stress in *B. cinerea*, which responds by forming chlamydospores, secreting oxylipins, and neutralizing fengycin. In turn, *Bacillus* has shifted its strategy, producing bacillaene to target fungal mitochondria. Additionally, *Bacillus* utilizes chitosan from the fungal cell wall, resulting in ecological competition between the two organisms.

## Discussion

In this study, we demonstrated that the *B. subtilis* ECM, in addition to playing a structural role in the communication with other bacteria and plants^78,79^, is essential for fungal hyphal colonization and antagonism against pathogenic fungi. Our observations indicate that the ECM can play an offensive or defensive role in interspecies interactions, depending on the competitor. In interactions with *Pseudomonas*, defense seems to be the prevalent function^78^; however, in interactions with *B. cinerea,* the role of ECM is more offensive. We show that the ECM components TasA and fengycin are crucial for antagonizing *B. cinerea*, acting synergistically while playing complementary roles, with each modifying *B. cinerea* physiology differently and triggering distinct metabolic changes.

Our data revealed that the ECM, specifically TasA, not only facilitates physical attachment but also orchestrates metabolite-driven antagonistic mechanisms, positioning the *B. subtilis* ECM as an active mediator of microbial competition. We specifically propose that the ECM component TasA has two unprecedented functions: one in altering the physiology of *B. cinerea* hyphae and the other as a stabilizer and carrier of the bioactive and highly sensitive molecule bacillaene. TasA alone directly disrupts *Botrytis* physiology by disorganizing its β-glucan layer, resulting in cellular and morphological damage. Moreover, our results highlight the specificity of fungal defenses against *B. subtilis* lipopeptides. For example, surfactin has a lethal effect on *Aspergillus*, inducing chlamydospore formation; however, it does not trigger this response in *B. cinerea^18^*. Conversely, fengycin has no apparent effect on the lifestyle of *Aspergillus* but does induce chlamydospore formation in *B. cinerea*. These findings suggest that the effectiveness of *B. subtilis* lipopeptides may be highly dependent on the unique membrane composition of different fungal species.

Furthermore, our results illustrate the complexity of the *Bacillus*.J*Botrytis* interaction, which involves bidirectional damage and dynamic chemical interplay, challenging the notion of purely antagonistic interkingdom bacteria□fungi interactions. Although *B. subtilis* initially suppresses fungal growth, the *B. cinerea* population initiates a chemical defense response by producing oxylipins that target *B. subtilis* and restructuring its cell morphology through hyperbranching and chlamydospore formation to increase survival. Some oxylipins, such as SDA, have been previously identified as mediators of survival strategies produced by the eukaryote *C. elegans* coexisting with *Pseudomonas*^80^, suggesting a conserved resilience mechanism among eukaryotes. The accumulation of Lyso-PC and ROS in *Botrytis* in response to TasA further indicates the complex biochemical interaction and metabolomic modulation of *B. cinerea* by *B. subtilis* ECM. Lyso-PC is implicated in oxidative stress in various organisms, and its accumulation in *B. cinerea*, alongside ROS, suggests the existence of a defense mechanism activated by the fungal cell in response to *B. cinerea* ECM. As part of this defense, *B. cinerea* partially neutralizes *B. subtilis* action by degrading fengycin, as previously reported for bacteria□bacteria interactions^81,82^ but not for interactions with fungi. Following the degradation of fengycin by *B. cinerea* to neutralize *B. subtilis* primary antifungal compounds, the production of another secondary metabolite from the *Bacillus* arsenal, bacillaene, increases. The role of bacillaene, which functions fungistatically rather than as a fungicide, exemplifies this adaptive strategy. Known to mediate microbe□microbe interactions^83^ and plant interactions, increased bacillaene production in response to fungal degradation of fengycin reflects the adaptability of *Bacillus* chemical weapons in response to *B. cinerea* counterattack. Additionally, we demonstrated that TasA binds to and stabilizes bacillaene, likely ensuring the fungistatic functionality. A previous study demonstrated the ability of bacillaene to bind the *E. coli* curli protein components CsgA and CsgB^84^, but in this case, it disrupted the assembly of amyloid fibers and thus biofilm formation, a strategy that confers *B. subtilis* with a competitive advantage in microbial interactions.

In the sequence of events leading to this interkingdom interaction, it may be predicted that following attachment to *B. cinerea* via TasA, *Bacillus* employs chitosanase to degrade chitosan from the *B. cinerea* cell wall and gain access to an extra carbon source. This ability to exploit *B. cinerea* for carbon is demonstrated by increased *B. subtilis* growth in coculture. Our results show that chitosanase acts synergistically with fengycin, increasing cell wall susceptibility to further fungal stress. Flow cytometry indicated that the presence of chitosan extends fengycin promoter activity, potentially stabilizing the antifungal arsenal of *B. subtilis*. Thus, the ECM, by supporting metabolite delivery and nutrient extraction, exemplifies an adaptive strategy in which *B. subtilis* can maintain *B. cinerea* in a state of fungistatic control without immediate eradication.

In short-term interactions, these adaptive responses may reflect a shift toward a stable antagonistic coexistence, consistent with ecological theories on microbial competition, suggesting that in some antagonistic interactions, competitors may reach a nonlethal equilibrium where metabolic adjustments allow for prolonged coculture^6,27^. This adaptability may represent a broader fungal strategy, wherein *B. cinerea* perceives specific ECM components from *B. subtilis*, thereby modulating its physiological defenses and enhancing survival. Competition upon initial contact can often evolve into a form of stable coexistence, where both *B. subtilis* and *B. cinerea* adapt and survive despite ongoing antagonism. Such coexistence may lead to spatial or resource partitioning, ultimately enabling both organisms to persist within a shared niche.

In conclusion, our data expand the understanding of microbial interactions by revealing the role of the bacterial ECM in mediating physical attachment and extracellular matrix-driven antagonism, with implications for microbial competition and survival strategies in resource-limited environments. Through such interactions, the *B. subtilis* ECM has emerged as a central element in microbial antagonism, shaping microbial interplay and sustaining *B. subtilis* survival under competitive conditions.

## Materials and Methods

### Strains, media and culture conditions

A complete list of the bacterial strains used in this study is shown in **Suppl. Table 3**. Bacterial cultures were grown in liquid lysogeny broth/Luria–Bertani (LB; 1% tryptone, 0.5% yeast extract and 0.5% NaCl) medium at 30°C (*Bacillus*) or 37°C (*E. coli*) with shaking on an orbital platform. The pH was adjusted to 7 prior to sterilization. When necessary, antibiotics were added to the media at appropriate concentrations. Isolate B05.10 of the necrotrophic fungus *B. cinerea* was cultured on potato dextrose agar (PDA, Oxoid) at 20°C under illumination with fluorescent light at a photofluency rate of 12□μmol□m^-2^□s^-1^ and a 12/12 h photoperiod. Conidia were harvested from the light-grown culture in sterile distilled water and filtered through a 40-μm cell strainer to remove the remaining hyphae. *Escherichia coli* DH5α was used for cloning and plasmid replication. *E. coli* BL21(DE3) and BL21(AI) were used for protein purification.

### B. subtilis mutants

*B. subtilis* mutants were generated using the SPP1 phage transduction method as previously described^85^. To generate double mutant strains, phage lysates were prepared from the ΔtasA and Δeps single mutant strains and subsequently transferred to a Δpps mutant. The mutants were verified through PCR and antibiotic resistance assays.

### Construction of fluorescence-labeled strains

The fluorescence labeling plasmid pKM008V was constructed for *Bacillus subtilis* strains. In brief, a 300 bp fragment of the Pveg promoter was extracted from pBS1C3 using restriction enzymes EcoRI and HindIII. The Pveg promoter was selected because of its role as a constitutive promoter in *B. subtilis.* After purification, the fragment was inserted into the pKM003 (YFP) or pDR183 (CFP) plasmid, which had been digested with the same enzymes. The plasmid was introduced into *B. subtilis* 168 by natural competence. Transformants were selected by plating on LB agar supplemented with antibiotics. To fluorescently label the extracellular matrix mutants, YFP or CFP was transferred from *B. subtilis* 168 using SPP1 phage transduction as previously described^85^.

### Bacterial**lJfungal** competition assay

*B. cinerea* was precultured by growing a germinated conidial suspension in potato dextrose broth (PDB, Oxoid) at 28°C for 24 h at 150 rpm. *B. subtilis* strains were grown on LB plates overnight at 28°C, and the resulting colonies were grown overnight in 5 mL of LB at 28°C on an orbital platform. Then, 10-15 germinated hyphae (microcolonies) were mixed with 200 µL of bacterial culture grown in 6-well plates with PDB at 28°C overnight at 150 rpm.

### Evaluation of *Bacillus* adhesion

To analyze the adhesion of different *Bacillus* ECM mutants to *Botrytis* hyphae, *Botrytis* was mixed with precultures of the different *Bacillus* strains constitutively labeled with YFP or CFP. After overnight incubation at 28°C in PDB, the *Botrytis* clumps were washed three times with water to remove unattached *Bacillus* cells. The washed hyphae were subsequently placed onto agarose-coated slides. For the *Bacillus* strains labeled with CFP, excitation and emission detection were performed at the corresponding wavelengths (excitation at 405 nm and emission detection between 450 and 550 nm), and likewise for YFP-labeled strains (excitation at 512 nm and emission detection between 520 and 600 nm). Images were obtained using a Leica Stellaris 8 confocal microscope with a 63x NA 1.3 Plan APO oil-immersion objective. For each experiment, the laser settings, scan speed, PMT or HyD detector gain, and pinhole aperture were kept constant across all acquired images. To count CFUs adhered to the fungus with each ECM mutant, the fungal mass was separated from the liquid bacterial culture by filtering with a 40 µm nylon filter to retain the fungus. The fungal mass was then washed once with Milli□Q (MQ) water to remove the free bacterial cells that were not attached to the fungi. The washed fungal mass was collected in a microtube and weighed to normalize the results to the mass of fungus. A pestle was then used to break the fungal structures and release the bacteria adhering to the fungus. The content was resuspended, and serial dilutions were performed. Finally, CFUs were counted and normalized to the fungal mass.

### Transmission electron microscopy

*B. cinerea* samples were fixed in 2.5% (v/v) glutaraldehyde and 4% (v/v) paraformaldehyde overnight at 4°C. After three washes in fixation mix, the samples were postfixed with 1% osmium tetroxide solution for 90 min at room temperature, followed by two washes and 15 min of stepwise dehydration in an ethanol series (30%, 50%, 70%, 90%, and 100% twice). Between the 50% and 70% steps, the samples were incubated en bloc in 2% uranyl acetate solution in 50% ethanol at 4°C overnight. Following dehydration, the samples were gradually embedded in low-viscosity Spurr’s resin as follows: resin:ethanol, 1:1, 4 h; resin:ethanol, 3:1, 4 h; and pure resin, overnight. The sample blocks were embedded in capsule molds containing pure resin for 72 h at 70°C. The samples were left to dry and visualized under an FEI TALOS F200X.

### Confocal laser scanning microscopy

Bacterial cell death within the colonies was assessed utilizing the LIVE/DEAD BacLight Bacterial Viability Kit (Invitrogen). The kit components were combined in equal proportions, and 2 μl of this mixture was used to stain 1 ml of the corresponding bacterial suspension. To visualize live or dead bacteria in the samples, sequential acquisitions were performed. Live bacteria were imaged using excitation at 488 nm, and emission was recorded between 499 and 554 nm, whereas dead bacteria were imaged through a subsequent acquisition via excitation at 561 nm and emission recorded between 592 and 688 nm.

Intracellular ROS levels were detected by staining with dihydrorhodamine 123 (DHR123; Sigma). Following incubation, DHR123 was added to the cell suspension to obtain a final concentration of 2 μg/mL, and the suspension was subsequently incubated for 5 min at room temperature. The cells were counterstained with the lipophilic dye FM4-64 (Thermo Fisher) to stain the plasma membrane. Images were obtained using a Leica Stellaris 8 confocal microscope with a 63x NA 1.3 Plan APO oil-immersion objective, with excitation at 488□nm and emission detection between 510 and 580 nm (for DHR123 fluorescence emission) and between 670 and 850 nm (for FM4-64 fluorescence emission). Image processing was performed using FIJI/ImageJ software^86^. For each experiment, the laser settings, scan speed, PMT or HyD detector gain, and pinhole aperture were kept constant across all acquired images.

### Protein purification

Recombinant His6-tagged TasA was purified as previously reported^87^. In brief, BL21 (DE3) *E. coli* was transformed with the pET22b-tasA plasmid. The resulting colonies were grown in 5 mL of LB medium at 37°C overnight at 150 rpm, reinoculated at a ratio of 1:100 in fresh LB medium and incubated at 37°C with shaking until an OD_600 of_ 0.7–0.8 was reached. At that point, protein expression was induced with 1 mM isopropyl β-D-1-thiogalactopyranoside (IPTG), and the mixture was incubated overnight at 28°C with shaking to induce inclusion body formation. The next day, the cells were harvested via centrifugation (7000 × g, 30 min, 4°C) and stored at -80°C until purification. After thawing, the pellet was resuspended in buffer A (Tris 50 mM, 150 mM NaCl; pH 8) supplemented with 0.2 mg/mL lysozyme, 1 mM PMSF, and 10x Cell Lysis Reagent (Sigma) and incubated for 1 h at 37°C. Then, 6 M GuHCl was added until saturation, and the mixture was incubated at 60°C overnight until complete solubilization occurred. The lysate was sonicated on ice (3 × 60 s, 60% amplitude) and centrifuged (110,000 × g, 1 h, 16°C). The resulting supernatant was passed through a 0.45-μm filter prior to affinity chromatography. The protein was purified using an AKTA Start FPLC system (GE Healthcare). The suspension was loaded into a HisTrap HP 5 mL column (GE Healthcare) previously equilibrated with binding buffer (50 mM Tris, 0.5 M NaCl, 20 mM imidazole, and 8 M urea; pH 8). Next, the protein was eluted from the column with elution buffer (50 mM Tris, 0.5 M NaCl, 500 mM imidazole, and 8 M urea; pH 8). Afterward, the purified protein was loaded into a HiPrep 26/10 desalting column (GE Healthcare), and the buffer was exchanged with 20 mM Tris or 50 mM NaCl to perform the corresponding experiments.

To purify chitosanase, recombinant His6-tagged chitosanase (Csn) was purified following a modified protocol for recombinant protein expression. Briefly, *E. coli* cultures carrying the Csn expression plasmid were grown in 250 mL of LB medium supplemented with ampicillin and incubated with shaking until an OD_600_ of 0.7 was reached. Expression was induced with IPTG, and the cultures were incubated for the optimal expression time before the cells were harvested by centrifugation (10,000 × g, 5 min). The cell pellets were lysed in washing buffer containing 50 mM Na_3_PO_4_ (pH 8), 500 mM NaCl, 10 mM imidazole, 1 mM PMSF, 0.2 mg/mL lysozyme, and 10x CelLytic® (Sigma□Aldrich) and incubated with shaking at room temperature for 30 min. The cells were then disrupted by sonication on ice (3 × 60 s pulses), and the lysates were clarified by centrifugation and filtered.

The lysate was loaded onto a HisTrap HP column (GE Healthcare) on an AKTA Start FPLC system equilibrated with washing buffer. The protein was eluted via elution buffer (20 mM Na*3*PO4, 500 mM NaCl, and 500 mM imidazole; pH 8). The purified Csn protein was dialyzed against 50 mM Tris-HCl and 50 mM NaCl (pH 7.4) using a HiPrep™ 26/10 Desalting column (GE Healthcare) and stored at -80°C until further use.

### EPS extraction and purification

The *B. subtilis* strain lacking *tasA* was used for EPS production using an overnight preculture of Ty liquid medium, and one milliliter was inoculated into 200 mL of Ty liquid medium for static culture in a 24-well microplate and left at 30°C for 5 days. The cells were harvested by centrifugation at 8000 × g for 10 min and washed with distilled water. After centrifugation, the cell pellets were resuspended in PBS and mildly sonicated to separate EPSs from the cells. The soluble fractions were stirred on ice, and a 100% TCA solution was added to precipitate proteins from the EPS extract. The mixture was left at 4°C overnight, and the proteins were removed by centrifugation at 10000 × g for 20 min. EPSs from the supernatant were precipitated by adding 5 volumes of chilled ethanol under continuous stirring and incubating at 4°C overnight. The crude EPS mixture was collected by centrifugation (10,000 × g, 20 min). The pellets were dialyzed against distilled water overnight (Spectra/Por^®^ dialysis membrane with a molecular weight cutoff of 3.5 kDa), and the resulting solution was filtered and applied to a size exclusion chromatography column (Hiprep^TM^ 16/60 Sephacryl^TM^ S-300HR), eluting with MQ water. Aliquots of each fraction were analyzed by the phenol sulfuric acid method, and the absorption at 490 nm was monitored in a multiplate reader (FLUOstar Omega reader, BMG LabTech). Fractions containing carbohydrates were pooled and lyophilized for further analysis.

### Immunolocalization of TasA by CLSM

To assess the binding of purified tagged TasA to *Botrytis* hyphae, we added 3 μM purified TasA and incubated it overnight. Next, the samples were applied to well slides treated with 0.1% poly-L-lysine (Sigma□Aldrich) and incubated for 2 h. After the samples were removed, they were fixed with fixation buffer (3% paraformaldehyde and 0.1% glutaraldehyde diluted in PBS) for 10 min. The wells were subsequently rinsed twice with PBS and incubated for 1 h in blocking buffer (3% w/v bovine serum albumin (BSA) and 0.2% v/v Triton X-100 in PBS). Following removal of the buffer, the wells were treated with the primary antibody (anti-His) at a concentration of 1:100 diluted in blocking buffer and incubated for 3 h. The wells were then washed three times with washing buffer (0.2% w/v BSA and 0.05% v/v Triton X-100 in PBS), with each incubation lasting for 5 min. The wells were subsequently incubated for 2 h with the secondary antibody goat anti-rabbit IgG-Atto488 at a dilution of 1:200 in blocking buffer. The samples were washed once with washing buffer and twice with PBS, with each incubation lasting for 5 min. The immunostained samples were fixed for 5 min with fixation buffer, followed by rinsing with PBS three times. As a negative control, immunostaining was performed without incubation with the primary antibody. Visualization of immunostaining was carried out using confocal laser scanning microscopy. For Atto-488 fluorescence, an excitation wavelength of 488 nm and emission wavelengths between 497 and 572 nm were employed. The Alexa Fluor 647 signal was visualized with an excitation wavelength of 561 nm, and emission was detected between 576 and 686 nm.

### TasA immunolabeling assay by TEM

To identify the specific localization of TasA on the cell wall of *Botrytis*, we first added 3 μM purified TasA to a *Botrytis* culture and incubated it overnight. Next, the cells were fixed and embedded as described in the TEM section. For immunolabeling assays, carbon-coated copper grids were deposited over the samples of *Botrytis*. After 2 h of incubation, the grids were washed in PBS for 5 min and blocked with Pierce protein-free (TBS) blocking buffer (Thermo Fisher) for 30 min. An anti-TasA primary antibody was used at a 1:150 dilution in blocking buffer, and the grids were deposited over the drops of antibody solution and incubated for 1 h at room temperature. The samples were washed three times with TBS-T (50 mM Tris-HCl, 150 mM NaCl, pH 7.5, and 0.1% Tween 20) for 5 min and then exposed to a 10-nm-diameter immunogold-conjugated secondary antibody (10 nm goat anti-rabbit conjugate, BBI solutions) for 1 h at a 1:50 dilution. The samples were then washed twice with TBS-T and once with water for 5 min each time. Finally, the grids were treated with 2% glutaraldehyde for 10 min, washed in water for 5 min, negatively stained with 1% uranyl acetate for 15 s and washed once with water for 15 s. The samples were left to dry, and images were acquired via a JEOL JEM-1400 transmission electron microscope at an accelerating voltage of 80 kV.

### MRI assays

*B. cinerea* cells (treated with purified TasA or protein buffer) were incubated for 24 h in PDB. Following incubation, all samples were embedded in 1.5% agar in Falcon tubes to ensure stable positioning for imaging. MRI experiments were conducted using a 9.4 T Bruker Biospec system equipped with 400 mT/m gradients and an Avance III console (Bruker BioSpin, Ettlingen, Germany). High-resolution T2-weighted images were acquired using a turbo-RARE sequence with the following parameters: TE = 33 ms, TR = 500 ms, 2 averages, a field of view (FOV) of 3.2 cm, a matrix size of 384 × 384, an in-plane resolution of 78 μm, and a slice thickness of 1 mm. These settings enabled detailed visualization of structural and texture variations in the *Botrytis* samples in response to TasA treatment.

### Plant infection assays

Assays of *B. cinerea* infection were carried out in 5–6-week-old plants. Fungal conidia were collected from cultures grown under illumination in sterile distilled water and filtered through a 40 μm cell strainer to remove the remaining hyphae. For inoculation, the conidial suspension was adjusted to 10^5^ conidia/mL in grape juice (100% pure organic). Each leaf was inoculated with 5 μL droplets of conidial suspension. The pots were covered with a plastic dome and placed in a growth chamber. After 24 h, the plastic dome was temporarily removed to apply the treatment (TasA) or buffer to the control plants. The leaves were imaged 72 h after inoculation, and the size of the lesions was determined using ImageJ software.

### RNA isolation and sequencing

RNA from *B. cinerea* was extracted from clumps disrupted with liquid nitrogen. After disruption, the suspensions of the pellets were resuspended in TRIzol reagent (Invitrogen). Total RNA extraction was then performed as indicated by the manufacturer. DNA removal was carried out by treatment with Nucleo-Spin RNA Plant (Macherey–Nagel). The integrity and quality of the total RNA were assessed with an Agilent 2100 Bioanalyzer (Agilent Technologies) via electrophoresis. The removal of rRNA was performed using the RiboZero rRNA Removal (Bacteria) Kit from Illumina, and 100-bp single-end read libraries were prepared via the TruSeq Stranded Total RNA Kit (Illumina). The libraries were sequenced using a NextSeq550 sequencer (Illumina). The raw reads were preprocessed with SeqTrimNext^88^ using specific next-generation sequencing (NGS) parameters. This preprocessing removed low-quality, ambiguous and low-complexity stretches; linkers; adapters; vector fragments; and contaminated sequences while keeping the longest informative parts of the reads. SeqTrimNext also discarded sequences less than 25□bp in length. The clean reads were subsequently aligned and annotated using the Bsub reference genome with Bowtie2^89^ in BAM files, which were then sorted and indexed using SAMtools v1.484^90^. Uniquely localized reads were used to calculate the read number value for each gene via Sam2counts (https://github.com/vsbuffalo/sam2counts). Differentially expressed genes (DEGs) were analyzed via DEgenes Hunter, which provides a combined p value calculated (based on Fisher’s method) using the nominal p values provided by edgeR^91^ and DEseq2. This combined p value was adjusted via the Benjamini□Hochberg (BH) procedure (false discovery rate approach) and used to rank all the obtained DEGs. For each gene, a combined p value < 0.05 and log2-fold change > 1 or < −1 were considered the significance thresholds.

### Metabolite extraction from liquid culture

The cultures were subsequently centrifuged to separate the cell and supernatant fractions. For the extraction of metabolites from the cell fraction, 1 mL of 80% methanol was added, and a tissue-lyser was used for 10 min to disrupt the cells. The mixture was then centrifuged at maximum speed, and the supernatant was transferred to a new tube. The methanol was evaporated using a speed vacuum concentrator and stored at -20°C until analysis by LC□MS. For the supernatant fraction, ethyl acetate was added at a 1:1 ratio, by volume, with the supernatant. The mixture was vortexed and rotated for 30 min. Subsequently, 4 mL of the upper phase was transferred to a new tube. The supernatant was evaporated using a speed vacuum concentrator and stored at -20°C until further analysis via LC□MS.

### Liquid chromatography**lJtandem** mass spectrometry (LC**lJMSlJMSlJMS**)

Nontarget metabolomics was performed by liquid chromatography□tandem mass spectrometry (LC□MS□MS) with a UHPLC coupled to a Q Exactive HF mass spectrometer as previously described^92^. In brief, UHPLC separation was performed using a C18 core□shell column (Kinetex, 50 × 1 mm, 1.7 µm particle size, 100 A pore size; Phenomenex, Torrance, USA). The mobile phases used were solvent (A), containing H2O (LC/MS grade, Fisher Scientific) + 0.1% formic acid (FA), and solvent (B), containing acetonitrile (LC/MS grade, Fisher Scientific) + 0.1% FA. After sample injection, a linear gradient of 5 min was used for the elution of small molecules, and the flow rate was set to 150 µl/min (microflow). The following separation conditions were used: 0–4 min from 5% to 50% solvent (B), 4–5 min from 50 to 99% B, followed by a 2 min washout phase at 99% B and a 3 min re-equilibration phase at 5% B. The measurements were conducted in positive mode, and the HESI parameters included a sheath gas flow rate of 30 L/min, an auxiliary gas flow rate of 10 L/min, and a sweep gas flow rate of 2 L/min. The spray voltage was set to 3.50 kV, the inlet capillary temperature was 250°C, the S-lens RF level was 50 V, and the auxiliary gas heater temperature was 200°C. The full MS survey scan acquisition range was set to 120–1,800 m/z with a resolution of 45,000, automatic gain control (AGC) of 1E6, and a maximum injection time of 100 ms with one microscan. DDA MS/MS spectra acquisition was performed in data-dependent acquisition (DDA) mode with TopN set to 5; as a consequence, the five most abundant precursor ions of the survey MS scan were subjected to MS/MS fragmentation. The resolution of the MS/MS spectra was set to 15,000, the AGC target was 5E5, and the maximum injection time was 50 ms. The quadrupole precursor selection width was set to 1 m/z. Normalized collision energy was applied stepwise at 25, 35, and 45°C. MS/MS scans were triggered in apex mode within 2–15 s from their first occurrence in a survey scan. Dynamic precursor exclusion was set to 5 s.

### Feature-based molecular networking and spectral library search

After LC□MS/MS acquisition, the raw spectra were converted to .mzML files using MSconvert (ProteoWizard). MS1 and MS/MS feature extraction was performed with Mzmine3^93^. For MS1 spectra, an intensity threshold of 1E5 was used, and for MS/MS spectra, an intensity threshold of 1E3 was used. For MS1 chromatogram building, a 10-ppm mass accuracy and a minimum peak intensity of 5E5 were set. Extracted ion chromatograms (XICs) were deconvolved using the baseline cutoff algorithm at an intensity of 1E5. After chromatographic deconvolution, the XICs were matched to the MS/MS spectra within 0.02 m/z and 0.2-minute retention time windows. Isotope peaks were grouped, and features from different samples were aligned with 10 ppm mass tolerance and 0.1-minute retention time tolerance. MS1 features without MS2 features assigned were filtered out of the resulting matrix, as were features that did not contain isotope peaks and that did not occur in at least three samples. After filtering, the gaps in the feature matrix were filled with a relaxed retention time tolerance of 0.2 min and a 10 ppm mass tolerance. Finally, the feature table was exported as a .csv file, and the corresponding MS/MS spectra were exported as .mgf files. Contaminant features observed in blank samples were filtered, and only those with a relative abundance ratio of blank to average lower than 30% were considered for further analysis. For feature-based molecular networking and spectrum library matching, the .mgf file was uploaded to GNPS^94,95^.

For molecular networking, the minimum cosine score was set to 0.7. The precursor ion mass tolerance was set to 0.01 Da, and the fragment ion mass tolerance was set to 0.01 Da. The minimum number of matched fragment peaks was set to 6, the minimum cluster size was set to 1 (MS cluster-off), and the library search minimum number of matched fragment peaks was set to 5. When analog searches were performed, the cosine score threshold was 0.7, and the maximum analog search mass difference was 100 m/z. Molecular networks were visualized with Cytoscape version 3.9.1^96^.

To enhance the chemical structural information in the molecular network, the generated .mgf file from MzMine3 was placed into Sirius 4 for chemical class and structure prediction^97^. The input data were automatically compared and classified against databases present in SIRIUS (Bio Database, GNPS, Natural Products, PubChem, PubMed). For molecular formula identification, the MS2 mass accuracy was set to 3 ppm. Chemical class annotations were performed with CSI: FingerID^98^ and CANOPUS^99,100^. Mirror plots were generated using GNPS and https://metabolomics-usi.ucsd.edu/, and the mzspec values of the selected features and the metabolites recorded in the MS/MS databases were compared (**Extended Data Fig. 9**). Annotations were performed according to the guidelines in ref.^101^ (**Suppl. Table 4**). Statistical analysis of the metabolomic datasets was performed via MetaboAnalyst v.5.0 after data were filtered by the interquartile range (IQR)^102^.

An automatic workflow for the analysis of the cell and supernatant fractions of *B. cinerea* and *B. subtilis* during coculture and *B. cinerea* treatment with TasA and fengycin at 6, 24 and 48 h (MSV000089552) can be accessed at https://massive.ucsd.edu/ProteoSAFe/dataset.jsp?task=b7579a1d14df42a09d30bd1418852db2 (feature-based molecular networking of the cell fraction:https://gnps.ucsd.edu/ProteoSAFe/status.jsp?task=4e4ade42ae60484886df1295747c5c71; feature-based molecular networking of the supernatant fraction: https://gnps.ucsd.edu/ProteoSAFe/status.jsp?task=c03281452dd342ccaf6962a051debb40).

### Native metabolomics

For the native metabolomics experiments, the same chromatographic parameters were used. In addition, 150 μL/min ammonium acetate buffer was infused postcolumn through a makeup pump and a PEEKT splitter, and a 3 μM TasA protein solution was infused with a 2 μL/min flow rate via the integrated syringe pump. The ESI settings were as follows: sheath gas flow, 40 arbitrary units; auxiliary gas flow, 10 arbitrary units; and sweep gas flow, 0 arbitrary units. The auxiliary gas temperature was set to 150°C. The spray voltage was set to 3 kV, and the inlet capillary was heated to 253°C. The S-lens level was set to 30 V. The MS scan range was set to 2500–4000 m/z with a resolution R_m/z_ 200 of 140,000 to 120,000 with 2 microscans. MS acquisition was performed in all-ion fragmentation (AIF) mode with R_m/z_ 200 with 20% HCD collision energy and an isolation window of 2500–4000 m/z. For native LC□MS data analysis, raw ion spectra were analyzed with Xcalibur software (Thermo Scientific) to identify changes in protein mass. Next, when a suspected ligand was found, feature tables from both the intact protein mass and the metabolomics data were matched by their retention time and a m/z offset corresponding to the masses of bacillaene B and dihydrobacillaene B.

### Minimum inhibitory concentration (MIC) assays

MIC assays were conducted in liquid LB medium using the twofold serial dilution method as outlined by the guidelines of the Clinical and Laboratory Standards Institute (2003). The highest concentration tested for the compound SDA was 1000 μg/mL. All experiments were performed in triplicate, and the MIC was determined as the lowest antibiotic concentration that inhibited growth by more than 90%.

### Polysaccharide affinity assay

These assays were conducted following established protocols^103,104^, employing 30 µg/mL purified TasA. In brief, TasA monomers were incubated with 3 mg of chitin beads (New England Biolabs), crab shell chitin, chitosan, β-glucan, cellulose (all from Sigma) or xylan (TCI America) in 800 mL of water. Following gentle agitation at 4°C overnight, the insoluble fraction was pelleted by centrifugation (13000 × g for 5 min), and the supernatant was collected. The insoluble fraction was washed three times with water and subsequently boiled in 1% SDS solution. The presence of protein in both the supernatant and pellet fractions was assessed via Tricine SDS□PAGE followed by Coomassie Brilliant Blue staining.

### Bacillaene extraction and purification

The extraction and purification of bacillaene were performed as previously described with some modifications^105,106^. In brief, cultures of *B. subtilis* (or Δpks as a control) were grown in 1 L batches in LB broth and incubated overnight at 28°C with shaking. Following incubation, the cells were removed by centrifugation (9 000 rpm, 10 min, 16°C), and the resulting cell-free supernatant was retained. Next, 250 mL of ethyl acetate was added for bacillaene extraction, and the mixture was incubated with agitation for 2 h and then left for 30 min to allow phase separation. The organic phase containing bacillaene was recovered and evaporated by lyophilization. The residue was dissolved in methanol and filtered through 0.45 µm filters (Econofltr PTFE). The filtered sample was chromatographed using a preparative HPLC column (Eclipse XDB-C18 5 μm, 9.4x250 mm). The mobile phases used were 20 mM NaPi (A) and acetonitrile (B), with the following elution profile: 0–2 min at 35% B, 2–8 min from 35–40% B, 8–10 min at 40% B, 10–12 min from 40–35% B, and 12–15 min at 35% B. The flow rate of the mobile phase was set to 1 mL/min, and the absorbance was monitored at 362 nm using a photodiode array (PDA). The presence of bacillaene was confirmed by comparing the chromatograms of the wild-type extract with those of the Δpks mutant extract, which lacked bacillaene, and by HPLC-MS by the MEDINA Foundation. The samples were analyzed using an Agilent 1200 Rapid Resolution HPLC connected to a Bruker maXis mass spectrometer. For separation, a Zorbax SB-C8 column (2.1 × 30 mm, 3.5 µm particle size) was used. The mobile phase consisted of two solvents: a 90:10 aqueous solution (solvent A) and a 10:90 aqueous solution (solvent B) of 13 mM ammonium formate and 0.01% TFA. The mass spectrometer was set to positive electrospray ionization (ESI) mode, with the following parameters: 4 kV capillary voltage, 11 L/min at 200°C for the drying gas flow, and 2.8 bar pressure in the nebulizer. The instrument was calibrated prior to sample injection via the ion cluster generated by TFA in the presence of Na+ ions. Each injected sample was subsequently recalibrated by infusing the TFA-Na calibrant before the chromatographic front appeared. Each chromatographic run was processed using Bruker’s internal algorithm for component extraction, and the most intense peaks, both by TIC in positive mode and absorbance at 210 nm, were considered for accurate mass interpretation and molecular formula determination. The combination of retention time and exact mass was used as a search criterion in the high-resolution mass spectrometry database of the MEDINA Foundation.

### Docking

Automated tertiary structure modeling of the elongation factor (EF) protein from *B. cinerea* B05.10 (Acc. no. XP_001560460.1), derived from the Bchbs1 and Bcmef1 genes, were generated via AlphaFold^107^. To identify potential binding sites for bacillaene (PubChem ID: 25144999) on the EF proteins, we utilized the web-based SwissDock program [www.swissdock.ch/docking]^108^ for automated molecular docking and thermodynamic analysis. SwissDock employs the EADock DSS algorithm^109^ to predict molecular interactions between a target protein and a small molecule. Docking was carried out using the “Accurate” setting with default parameters and no predefined region of interest (blind docking). Binding energies were calculated using CHARMM (Chemistry at HARvard Macromolecular Mechanics), integrated within SwissDock software, and the most favorable energies were assessed via fast analytical continuum treatment of solvation (FACTS). The energy results were then scored and ranked on the basis of full fitness (kcal mol−1), with spontaneous binding indicated by the estimated Gibbs free energy ΔG (kcal mol−1). The negative ΔG values suggest that the binding process is highly spontaneous. Visualization of the modeling and docking results was performed using UCSF Chimera v1.8 software.

### Flow cytometry assays

The cells were grown in 24-well plates filled with 1 mL of MSgg at 28□°C. At different time points, the pellicle was carefully removed, and the cells from the spent medium were recovered in 500□μL of PBS after centrifugation (14,000 rpm, 3 min) and resuspended with a 25G needle. The cells were gently sonicated (12 pulses of 5 s and 30% amplitude) to ensure complete resuspension, fixed in 4% paraformaldehyde in PBS and washed three times in PBS. The flow cytometry runs were performed with 200□μl cell suspensions in 800□μl of GTE buffer (50□mM glucose, 10□mM EDTA, 20□mM Tris-HCl; pH 8), and the cells were quantified on a Beckman Coulter Gallios™ flow cytometer using 488□nm excitation. YFP fluorescence was detected with a 525/40 BP filter. The data were collected using Gallios™ Software v1.2 and further analyzed using Flowing Software v2.5.1. Unlabeled cells (*B. subtilis* wild type) were used as a negative control for promoter expression analysis.

### Chitosan level quantification

Changes in the chitosan composition were determined via eosin Y labeling. The samples were subsequently resuspended in citrate-phosphate buffer (0.2 M NaH_2_PO_4_, 0.1 M K citrate; pH 6) with 1 μg/ml eosin Y (final concentration). After 10 min of incubation at room temperature, the cells were washed two times with citrate-phosphate buffer and placed on 1% agarose pads. Finally, the stained cells were imaged with an excitation wavelength of 488 nm, and emission was detected between 510 and 640 nm. Images were obtained using a Leica Stellaris 8 confocal microscope with a 63x NA 1.3 Plan APO oil-immersion objective. Processing and signal intensity measurement were performed via FIJI/ImageJ. For each experiment, the laser settings, scan speed, HyD detector gain, and pinhole aperture were kept constant across all acquired images.

## Data availability

All the raw RNA-seq data have been submitted to the Gene Expression Omnibus (GEO) and can be accessed through GEO series accession no. GSE287578 (URL: https://www.ncbi.nlm.nih.gov/geo/query/acc.cgi?acc=GSE287578. All the metabolomics data are deposited at https://massive.ucsd.edu/with the identifier MSV000089552.

## Acknowledgments

We thank Saray Morales for providing technical support. We also thank Alicia Esteban from the IHSM microscopy unit, and David Navas, Cristina Lucena and Adolfo Martínez from the SCAI microscopy until, for their technical support in confocal and electronic microscopy. We also thank Josefa Gómez Maldonado and Luís Díaz from the Ultrasequencing Unit of the SCBI-UMA for RNA sequencing and bioinformatic analysis and Mercedes Martín Rufián and Casimiro Cárdenas García from the Proteomic Unit of the SCAI-UMA for their technical suggestions, protein sequencing, and LC□MS analysis. This work was supported by grants from an ERC Starting Grant (BacBio 637971), Agencia Estatal de Investigación of the Ministerio de Ciencia e Innovación (PID2019-107724GB-I00 and PID2022-141664NB-I00), and Junta de Andalucía (P20_00479). A.I.P.L. is funded by the program FPU (FPU19/00289) and the program Plan Propio de Investigación y Transferencia from Universidad de Málaga. C.M.S. is funded by grants from Consolidación Investigadora (CSN2022-135744) and Proyectos dirigidos por jóvenes investigadores de la Universidad de Málaga (B1-2021_21). D.V.C. is funded by the program Incorporación de Doctores PAIDI from Junta de Andalucía (DOC_00266) and Proyecto Jóvenes Investigadores from the Plan Propio de Universidad de Málaga (B1-2021_34). P.S. was supported by the European Union’s Horizon Europe Research and Innovation Programme through a Marie Skłodowska-Curie fellowship (no. 101108450 MeStaLeM). D.P. was supported by the German Research Foundation (DFG) through the Cluster of Excellence Controlling Microbes to Fight Infections, CMFI (EXC2124).

---

**Extended Data Figure 1. Enrichment analysis of differentially expressed *B. cinerea* genes during interaction with *B. subtilis.* A)** KEGG pathway enrichment analysis showing the deregulation of pathways related to glutathione metabolism, secondary metabolite biosynthesis, and phospholipid metabolism after six hours of coculture. **B)** Gene Ontology (GO) terms enriched in genes upregulated in *B. cinerea* after six hours of coculture. **C)** GO terms enriched by genes downregulated in B. cinerea after six hours of coculture.

**Extended Data Figure 2. Enrichment analysis of differentially expressed *B. subtilis* genes during interaction with *B. cinerea*.** KEGG pathway enrichment analysis showing the deregulation of pathways related to the biosynthesis of secondary metabolites.

**Extended Data Figure. 3. Metabolomic analysis revealed differential accumulation of metabolite classes during the *Bacillus-Botrytis* interaction. A)** Top 25 features with the highest median weighted sum of absolute regression coefficient scores determined by PLS-DA, calculated using MetaboAnalyst. These features were identified as key metabolites discriminating cocultures from single cultures of each microbe after 24 h in the cell fraction. The feature IDs are annotated with their chemical classes via Classyfire classification. The left side of the figure indicates the microbial origin of the different accumulated metabolite classes. **B)** Heatmap of hierarchical clustering of the top 350 features within impacted molecular families during the coculture of *B. cinerea* and *B. subtilis* compared with monoculture of each after 24 h in the cell fraction. The color gradient in the heatmap indicates the relative fold change of each metabolite between the groups. The circular graphs show the percentage distribution of metabolite classes with altered accumulation, with the color code inside the donut chart representing each chemical class according to NPClassifier. **C)** Quantification of ROS levels in *B. cinerea* after treatment with 150 µM lysoPC or no treatment for 24 h. The whisker plot shows all the measurements (pink dots), medians (black line), and minimum and maximum values (whisker ends). For all experiments, the results from at least three biological replicates are shown. Statistical significance was assessed via a t test, with quadruple asterisks indicating significant differences at P < 0.0001.

**Extended Data Figure 4. A) Top:** Transmission electron micrograph of negatively stained thin sections of untreated *B. cinerea* hyphae with immunogold labeling. The scale bar represents 1 μm for the image on the left and 0.2 μm for the zoomed-in image on the right. **Bottom:** Immunocytochemistry of *Botrytis* hyphae treated with TasA without anti-TasA antibodies but with secondary antibodies as a control. Scale bars represent 20 µm. **B)** Coomassie gel of polysaccharide affinity assays with purified TasA. **C)** Enrichment analysis of differentially expressed *B. cinerea* genes after six hours of treatment with 3 µm purified TasA. GO terms enriched by upregulated (top) and downregulated (bottom) genes of *B. cinerea* after treatment with TasA. D) Quantification of tubulin accumulation in pear hyphae treated with 3 µm purified TasA or left untreated. The whisker plot shows all the measurements (pink dots), medians (black line), and minimum and maximum values (whisker ends). For all experiments, the results from at least three biological replicates are shown. Statistical significance was assessed via a t test, with quadruple asterisks indicating significant differences at P < 0.0001.

**Extended Data Figure 5. Differential effects of EPS and fengycin on *B. cinerea* physiology and stress responses. A)** Representative confocal microscopy images showing ROS accumulation and the fungal membrane (stained with FM4-64) in *B. cinerea* hyphae treated with EPS. Scale bars represent 20 µm. The lower panels show transmission micrographs of *B. cinerea* treated with 0.5 mg/mL purified EPS, revealing nonpronounced ultrastructural disorganization or autophagosome formation. Scale bars represent 2 μm. **B)** Quantification of ROS levels in *B. cinerea* without treatment, with H_2_O_2_ as positive control, after treatment with 3 µM TasA, 0.5 mg/mL EPS or 10 µM fengycin for 24 h. Whisker plot shows all measurements (pink dots), medians (black line), and minimum and maximum (whiskers ends). In all experiments, at least three biological replicates are shown. Statistical significance was assessed ANOVA by one-way ANOVA with Dunnett’s multiple comparisons test (each treatment vs BcB05 control), with quadruple asterisk indicating significant differences at P < 0.0001.

**Extended Data Figure 6. Differentially abundant metabolite accumulation in *B. cinerea* supernatants after TasA and fengycin treatments.** Volcano plots showing metabolites with significantly greater or lower accumulation in the *B. cinerea* supernatant after 24 h of treatment with TasA (left) or fengycin (right) than in the supernatant of untreated *B. cinerea*. Analyses were performed using MetaboAnalyst.

**Extended Data Figure. 7. KEGG pathway-based schematics of oxylipins accumulated in *Botrytis cinerea* supernatant after treatment with fengycin**. Two pathway diagrams representing the accumulation of oxylipins derived from linoleic (top scheme) and linolenic (bottom scheme) acids in the *B. cinerea* supernatant. The accumulated metabolites are colored according to the same scheme as the border node color in **Figure 4A**.

**Extended Data Figure 8. Molecular family analysis of bacillaene in *B. cinerea* supernatant after interaction with wild-type *B. subtilis* compared with *B. cinerea* interacting with** Δ**pps after 48h**. Pie charts represent the mean peak abundance of metabolites in the supernatant, with node shapes indicating the level of metabolite identification according to GNPS libraries.

**Extended Data Figure 9. Mirror plots comparing spectra from named features throughout the manuscript to standard spectra deposited in GNPS**. The upper part of the plot (black lines) represents the MS spectra of the candidate feature, and the lower part (green lines) represents the MS spectra of the standard compound. Mirror plots were generated using https://metabolomics-usi.ucsd.edu/.

**Supplementary Table 1. *B. cinerea* B05.10 putative secreted proteins upregulated upon challenging with Fengycin.**

**Supplementary Table 2. List of mutated genes in *Botrytis cinerea* treated with bacillaene for one week compared with the *Botrytis* control**

**Supplementary Table 3. List of strains used in this study.**

**Supplementary Table 4. Named features throughout the manuscript and their parameters for identification within the framework of Sumner *et al*., 2007.**

## References

1. Deveau, A. et al. Bacterial-fungal interactions: Ecology, mechanisms and challenges. FEMS Microbiol Rev 42, 335–352 (2018).

2. Michael E. Hibbing, Clay Fuqua, Matthew R. Parsek, and S. B. P. Bacterial competition: surviving and thriving in the microbial jungle. Nat Rev Microbiol 123, 75–88 (2015).

3. Cordovez, V., Dini-Andreote, F., Carrión, V. J. & Raaijmakers, J. M. Ecology and evolution of plant microbiomes. Annu Rev Microbiol 73, 69– 88 (2019).

4. Granato, E. T., Meiller-Legrand, T. A. & Foster, K. R. The Evolution and Ecology of Bacterial Warfare. Current Biology vol. 29 R521–R537 Preprint at 10.1016/j.cub.2019.04.024 (2019).

5. Foster, K. R. & Bell, T. Competition, not cooperation, dominates interactions among culturable microbial species. Current Biology 22, 1845–1850 (2012).

6. Peterson, S. B., Bertolli, S. K. & Mougous, J. D. The Central Role of Interbacterial Antagonism in Bacterial Life. Current Biology 30, R1203– R1214 (2020).

7. Ongena, M. & Jacques, P. Bacillus lipopeptides: versatile weapons for plant disease biocontrol. Trends in Microbiology vol. 16 115–125 Preprint at 10.1016/j.tim.2007.12.009 (2008).

8. Cazorla, F. M. et al. Isolation and characterization of antagonistic Bacillus subtilis strains from the avocado rhizoplane displaying biocontrol activity. J Appl Microbiol 103, 1950–1959 (2007).

9. Bouchard-Rochette, M. et al. Bacillus pumilus PTB180 and Bacillus subtilis PTB185: Production of lipopeptides, antifungal activity, and biocontrol ability against Botrytis cinerea. Biological Control 170, 104925 (2022).

10. Kovács, Á. T. Bacillus subtilis. Trends Microbiol 27, 724–725 (2019).

11. Kiesewalter, H. T. et al. Genomic and Chemical Diversity of Bacillus subtilis Secondary Metabolites against Plant Pathogenic Fungi. mSystems 6, (2021).

12. Romero, D., Aguilar, C., Losick, R. & Kolter, R. Amyloid fibers provide structural integrity to Bacillus subtilis biofilms. Proc Natl Acad Sci U S A 107, 2230–2234 (2010).

13. Romero, D., Vlamakis, H., Losick, R. & Koltera, R. Functional analysis of the accessory protein TapA in Bacillus subtilis amyloid fiber assembly. J Bacteriol 196, 1505–1513 (2014).

14. Branda, S. S., Chu, F., Kearns, D. B., Losick, R. & Kolter, R. A major protein component of the Bacillus subtilis biofilm matrix. Mol Microbiol 59, 1229–1238 (2006).

15. Berlanga-Clavero, M. V. et al. Bacillus subtilis biofilm matrix components target seed oil bodies to promote growth and anti-fungal resistance in melon. Nat Microbiol 7, 1001–1015 (2022).

16. Molina-Santiago et al. The extracellular matrix protects Bacillus subtilis colonies from Pseudomonas invasion and modulates plant co-colonization. Nat Commun 10, (2019).

17. Kjeldgaard, B., et al. Fungal hyphae colonization by Bacillus subtilis relies on biofilm matrix components. Biofilm 1, (2019).

18. Richter, A. et al. Enhanced surface colonisation and competition during bacterial adaptation to a fungus. Nat Commun 15, (2024).

19. Anckaert, A. et al. The biology and chemistry of a mutualism between a soil bacterium and a mycorrhizal fungus. Curr Biol (2024) doi:10.1016/j.cub.2024.09.019.

20. Wang, C. & Kuzyakov, Y. Mechanisms and implications of bacterial – fungal competition for soil resources. ISME J 18, (2024).

21. Bouchard-Rochette, M. et al. Bacillus pumilus PTB180 and Bacillus subtilis PTB185: Production of lipopeptides, antifungal activity, and biocontrol ability against Botrytis cinerea. Biological Control 170, 104925 (2022).

22. Samaras, A., Karaoglanidis, G. S. & Tzelepis, G. Insights into the multitrophic interactions between the biocontrol agent Bacillus subtilis MBI 600, the pathogen Botrytis cinerea and their plant host. Microbiol Res 248, 126752 (2021).

23. Bu, S. et al. Bacillus subtilis L1-21 as a biocontrol agent for postharvest gray mold of tomato caused by Botrytis cinerea. Biological Control 157, 104568 (2021).

24. Kodera, S. M., Das, P., Gilbert, J. A. & Lutz, H. L. iScience Conceptual strategies for characterizing interactions in microbial communities. doi:10.1016/j.isci.

25. Maynard Smith, J. The theory of games and the evolution of animal conflicts. J Theor Biol 47, 209–221 (1974).

26. Poullain, V., Gandon, S., Brockhurst, M. A., Buckling, A. & Hochberg, M. E. The evolution of specificity in evolving and coevolving antagonistic interactions between a bacteria and its phage. Evolution (N Y) 62, 1–11 (2008).

27. Ghoul, M. & Mitri, S. The Ecology and Evolution of Microbial Competition. Trends Microbiol 24, 833–845 (2016).

28. Kiesewalter, H. T. et al. Genomic and Chemical Diversity of Bacillus subtilis Secondary Metabolites against Plant Pathogenic Fungi. mSystems 6, (2021).

29. Touré, Y., Ongena, M., Jacques, P., Guiro, A. & Thonart, P. Role of lipopeptides produced by Bacillus subtilis GA1 in the reduction of grey mould disease caused by Botrytis cinerea on apple. J Appl Microbiol 96, 1151–1160 (2004).

30. Bouchard-Rochette, M. et al. Bacillus pumilus PTB180 and Bacillus subtilis PTB185: Production of lipopeptides, antifungal activity, and biocontrol ability against Botrytis cinerea. Biological Control 170, 104925 (2022).

31. Cámara-almirón, J., Navarro, Y. & Magno-pérez-bryan, M. C. Dual functionality of the TasA amyloid protein in Bacillus physiology and fitness on the phylloplane. bioRxiv 31, (2019).

32. Hussein, W. Fengycin or plipastatin? A confusing question in bacilli. Biotechnologia 100, 47–55 (2019).

33. Romero, D., Vlamakis, H., Losick, R. & Kolter, R. An accessory protein required for anchoring and assembly of amyloid fibres in B. subtilis biofilms. Mol Microbiol 80, 1155–1168 (2011).

34. Vinayavekhin, N., Mahipant, G., Vangnai, A. S. & Sangvanich, P. Untargeted metabolomics analysis revealed changes in the composition of glycerolipids and phospholipids in Bacillus subtilis under 1-butanol stress. Appl Microbiol Biotechnol 99, 5971–5983 (2015).

35. Willdigg, J. R. & Helmann, J. D. Mini Review: Bacterial Membrane Composition and Its Modulation in Response to Stress. Frontiers in Molecular Biosciences vol. 8 Preprint at 10.3389/fmolb.2021.634438 (2021).

36. Patton-Vogt, J. & de Kroon, A. I. P. M. Phospholipid turnover and acyl chain remodeling in the yeast ER. Biochimica et Biophysica Acta - Molecular and Cell Biology of Lipids vol. 1865 Preprint at 10.1016/j.bbalip.2019.05.006 (2020).

37. Henry, S. A., Kohlwein, S. D. & Carman, G. M. Metabolism and regulation of glycerolipids in the yeast Saccharomyces cerevisiae. Genetics 190, 317–349 (2012).

38. Xu, J. & Huang, X. Lipid Metabolism at Membrane Contacts: Dynamics and Functions Beyond Lipid Homeostasis. Frontiers in Cell and Developmental Biology vol. 8 Preprint at 10.3389/fcell.2020.615856 (2020).

39. Stålberg, K., Neal, A. C., Ronne, H. & Ståhl, U. Identification of a novel GPCAT activity and a new pathway for phosphatidylcholine biosynthesis in S. cerevisiae. J Lipid Res 49, 1794–1806 (2008).

40. Van Meer, G., Voelker, D. R. & Feigenson, G. W. Membrane lipids: Where they are and how they behave. Nature Reviews Molecular Cell Biology vol. 9 112–124 Preprint at 10.1038/nrm2330 (2008).

41. Wi, S. J., Seo, S. Y., Cho, K., Nam, M. H. & Park, K. Y. Lysophosphatidylcholine enhances susceptibility in signaling pathway against pathogen infection through biphasic production of reactive oxygen species and ethylene in tobacco plants. Phytochemistry 104, 48–59 (2014).

42. Schilling, T. & Eder, C. Importance of lipid rafts for lysophosphatidylcholine-induced caspase-1 activation and reactive oxygen species generation. Cell Immunol 265, 87–90 (2010).

43. Watanabe, N. et al. Activation of mitogen-activated protein kinases by lysophosphatidylcholine-induced mitochondrial reactive oxygen species generation in endothelial cells. American Journal of Pathology 168, 1737– 1748 (2006).

44. Yuan, Y., Schoenwaelder, S. M., Salem, H. H. & Jackson, S. P. The bioactive phospholipid, lysophosphatidylcholine, induces cellular effects via G-protein-dependent activation of adenylyl cyclase. Journal of Biological Chemistry 271, 27090–27098 (1996).

45. Barman, A., Gohain, D., Bora, U. & Tamuli, R. Phospholipases play multiple cellular roles including growth, stress tolerance, sexual development, and virulence in fungi. Microbiol Res 209, 55–69 (2018).

46. Dührkop, K. et al. Systematic classification of unknown metabolites using high-resolution fragmentation mass spectra. Nat Biotechnol 39, 462–471 (2021).

47. Lichius, A., Berepiki, A. & Read, N. D. Form follows function - The versatile fungal cytoskeleton. Fungal Biol 115, 518–540 (2011).

48. Francisco, C. S., Ma, X., Zwyssig, M. M., McDonald, B. A. & Palma-Guerrero, J. Morphological changes in response to environmental stresses in the fungal plant pathogen Zymoseptoria tritici. Sci Rep 9, 1–18 (2019).

49. Spraker, J. E., Sanchez, L. M., Lowe, T. M., Dorrestein, P. C. & Keller, N. P. Ralstonia solanacearum lipopeptide induces chlamydospore development in fungi and facilitates bacterial entry into fungal tissues. ISME Journal 10, 2317–2330 (2016).

50. Li, L. et al. Induction of chlamydospore formation in fusarium by cyclic lipopeptide antibiotics from bacillus subtilis C2. J Chem Ecol 38, 966–974 (2012).

51. Beccaccioli, M., Reverberi, M. & Scala, V. Fungal lipids: Biosynthesis and signalling during plant-pathogen interaction. Frontiers in Bioscience - Landmark 24, 172–185 (2019).

52. Beccaccioli, M. et al. Fungal and bacterial oxylipins are signals for intra- and inter-cellular communication within plant disease. Front Plant Sci 13, (2022).

53. Liu, H., Zhang, X., Chen, W. & Wang, C. The regulatory functions of oxylipins in fungi: A review. J Basic Microbiol 63, 1073–1084 (2023).

54. Tsitsigiannis, D. I. & Keller, N. P. Oxylipins as developmental and host-fungal communication signals. Trends Microbiol 15, 109–118 (2007).

55. Niu, M. et al. Fungal oxylipins direct programmed developmental switches in filamentous fungi. Nat Commun 11, 1–13 (2020).

56. Brodhun, F. & Feussner, I. Oxylipins in fungi. FEBS Journal 278, 1047– 1063 (2011).

57. Bielčik, M., Aguilar-Trigueros, C. A., Lakovic, M., Jeltsch, F. & Rillig, M. C. The role of active movement in fungal ecology and community assembly. Mov Ecol 7, 1–12 (2019).

58. Pakkir Shah, A. K., et al. Statistical analysis of feature-based molecular networking results from non-targeted metabolomics data. Nat Protoc (2024) doi:10.1038/s41596-024-01046-3.

59. Rigolet, A. et al. Lipopeptides as rhizosphere public goods for microbial cooperation. Microbiol Spectr 12, 1–13 (2024).

60. Vela-Corcia, D. et al. Cyclo(Pro-Tyr) elicits conserved cellular damage in fungi by targeting the [H+]ATPase Pma1 in plasma membrane domains. Commun Biol 7, 1253 (2024).

61. Hansen, M. L. et al. Resistance towards and biotransformation of a Pseudomonas-produced secondary metabolite during community invasion. ISME Journal 18, (2024).

62. Rigolet, A. et al. Lipopeptides as rhizosphere public goods for microbial cooperation. Microbiol Spectr 12, 1–13 (2024).

63. Miao, S., Liang, J., Xu, Y., Yu, G. & Shao, M. Bacillaene, sharp objects consist in the arsenal of antibiotics produced by Bacillus. J Cell Physiol 1995, (2023).

64. Kiesewalter, H. T. et al. Genomic and Chemical Diversity of Bacillus subtilis Secondary Metabolites against Plant Pathogenic Fungi. mSystems 6, (2021).

65. Li, H. et al. Bacillaenes: Decomposition Trigger Point and Biofilm Enhancement in Bacillus. ACS Omega 6, 1093–1098 (2021).

66. Li, H. et al. An Effective Strategy for Identification of Highly Unstable Bacillaenes. J Nat Prod 82, 3340–3346 (2019).

67. Patel, P. S. et al. Bacillaene, a novel inhibitor of procaryotic protein synthesis produced by Bacillus subtilis: production, taxonomy, isolation, physico-chemical characterization and biological activity. J Antibiot (Tokyo) 48, 997–1003 (1995).

68. Miao, S., Liang, J., Xu, Y., Yu, G. & Shao, M. Bacillaene, sharp objects consist in the arsenal of antibiotics produced by Bacillus. J Cell Physiol 1995, (2023).

69. Butcher, R. A. et al. The identification of bacillaene, the product of the PksX megacomplex in Bacillus subtilis. Proc Natl Acad Sci U S A 104, 1506–1509 (2007).

70. Molina-Santiago, C. et al. Chemical interplay and complementary adaptative strategies toggle bacterial antagonism and co-existence. Cell Rep 36, (2021).

71. Molina-Santiago, C. et al. Chemical interplay and complementary adaptative strategies toggle bacterial antagonism and co-existence. Cell Rep 36, (2021).

72. Marvasi, M., Visscher, P. T. & Casillas Martinez, L. Exopolymeric substances (EPS) from Bacillus subtilis: Polymers and genes encoding their synthesis. FEMS Microbiology Letters vol. 313 1–9 Preprint at 10.1111/j.1574-6968.2010.02085.x (2010).

73. Reher, R. et al. Native metabolomics identifies the rivulariapeptolide family of protease inhibitors. Nat Commun 13, 1–12 (2022).

74. Choi, Y. J., Kim, E. J., Piao, Z., Yun, Y. C. & Shin, Y. C. Purification and characterization of chitosanase from Bacillus sp. strain KCTC 0377BP and its application for the production of chitosan oligosaccharides. Appl Environ Microbiol 70, 4522–4531 (2004).

75. Su, P. C., Hsueh, W. C., Chang, W. S. & Chen, P. T. Enhancement of chitosanase secretion by Bacillus subtilis for production of chitosan oligosaccharides. J Taiwan Inst Chem Eng 79, 49–54 (2017).

76. Pechsrichuang, P. et al. Bioconversion of chitosan into chito-oligosaccharides (CHOS) using family 46 chitosanase from Bacillus subtilis (BsCsn46A). Carbohydr Polym 186, 420–428 (2018).

77. Baker, L. G., Specht, C. A., Donlin, M. J. & Lodge, J. K. Chitosan, the deacetylated form of chitin, is necessary for cell wall integrity in Cryptococcus neoformans. Eukaryot Cell 6, 855–867 (2007).

78. Molina-Santiago et al. The extracellular matrix protects Bacillus subtilis colonies from Pseudomonas invasion and modulates plant co-colonization. Nat Commun 10, (2019).

79. Berlanga-Clavero, M. V. et al. Bacillus subtilis biofilm matrix components target seed oil bodies to promote growth and anti-fungal resistance in melon. Nat Microbiol 7, 1001–1015 (2022).

80. Nandakumar, M. & Tan, M. W. Gamma-linolenic and stearidonic acids are required for basal immunity in Caenorhabditis elegans through their effects on p38 MAP kinase activity. PLoS Genet 4, (2008).

81. Hansen, M. L. et al. Resistance towards and biotransformation of a Pseudomonas-produced secondary metabolite during community invasion. ISME Journal 18, (2024).

82. Hoefler, B. C. et al. Enzymatic resistance to the lipopeptide surfactin as identified through imaging mass spectrometry of bacterial competition. Proc Natl Acad Sci U S A 109, 13082–13087 (2012).

83. Erega, A., et al. Bacillaene Mediates the Inhibitory Effect of Bacillus Subtilis on Campylobacter Jejuni Biofilms. 10 (2021).

84. Cordisco, E., Zanor, M. I., Moreno, D. M. & Serra, D. O. Selective inhibition of the amyloid matrix of Escherichia coli biofilms by a bifunctional microbial metabolite. NPJ Biofilms Microbiomes 9, (2023).

85. Thorne, C. B. & Mele, J. Prophage interference in Bacillus subtilis 168. Microb.Genet.Bull. **No.**36, 27–29 (1974).

86. Schindelin, J., et al. Fiji: An open-source platform for biological-image analysis. Nat Methods 9, 676–682 (2012).

87. Mammeri, N. El et al. Molecular architecture of bacterial amyloids in Bacillus biofilms. FASEB Journal 33, 12146–12163 (2019).

88. Falgueras, J. et al. SeqTrim: A high-throughput pipeline for pre-processing any type of sequence read. BMC Bioinformatics 11, (2010).

89. Langmead, B. & Salzberg, S. L. Fast gapped-read alignment with Bowtie 2. Nat Methods 9, 357–359 (2012).

90. Li, H. et al. The Sequence Alignment/Map format and SAMtools. Bioinformatics 25, 2078–2079 (2009).

91. Robinson, M. D., McCarthy, D. J. & Smyth, G. K. edgeR: A Bioconductor package for differential expression analysis of digital gene expression data. Bioinformatics 26, 139–140 (2009).

92. Stincone, P. et al. Evaluation of Data-Dependent MS/MS Acquisition Parameters for Non-Targeted Metabolomics and Molecular Networking of Environmental Samples: Focus on the Q Exactive Platform. Anal Chem 95, 12673–12682 (2023).

93. Schmid, R. et al. Integrative analysis of multimodal mass spectrometry data in MZmine 3. Nat Biotechnol 41, 447–449 (2023).

94. Wang, M. et al. Sharing and community curation of mass spectrometry data with Global Natural Products Social Molecular Networking. Nat Biotechnol 34, 828–837 (2016).

95. Petras, D. et al. GNPS Dashboard: collaborative exploration of mass spectrometry data in the web browser. Nat Methods 19, 134–136 (2022).

96. Paul Shannon, 1 et al. Cytoscape: A Software Environment for Integrated Models. Genome Res 13, 426 (1971).

97. Dührkop, K. et al. SIRIUS 4: a rapid tool for turning tandem mass spectra into metabolite structure information. Nat Methods 16, 299–302 (2019).

98. Dührkop, K., Shen, H., Meusel, M., Rousu, J. & Böcker, S. Searching molecular structure databases with tandem mass spectra using CSI:FingerID. Proc Natl Acad Sci U S A 112, 12580–12585 (2015).

99. Kim, H. W. et al. NPClassifier: A Deep Neural Network-Based Structural Classification Tool for Natural Products. J Nat Prod 84, 2795–2807 (2021).

100. Dührkop, K. et al. Systematic classification of unknown metabolites using high-resolution fragmentation mass spectra. Nat Biotechnol 39, 462–471 (2021).

101. Sumner, L. W. et al. Proposed minimum reporting standards for chemical analysis: Chemical Analysis Working Group (CAWG) Metabolomics Standards Initiative (MSI). Metabolomics 3, 211–221 (2007).

102. Pang, Z. et al. MetaboAnalyst 5.0: Narrowing the gap between raw spectra and functional insights. Nucleic Acids Res 49, W388–W396 (2021).

103. Mirzadi Gohari, A., et al. Effector discovery in the fungal wheat pathogen Zymoseptoria tritici. Mol Plant Pathol 16, 931–945 (2015).

104. Sánchez-Vallet, A. et al. A secreted LysM effector protects fungal hyphae through chitin-dependent homodimer polymerization. PLoS Pathog 16, 1– 21 (2020).

105. Cordisco, E., Zanor, M. I., Moreno, D. M. & Serra, D. O. Selective inhibition of the amyloid matrix of Escherichia coli biofilms by a bifunctional microbial metabolite. NPJ Biofilms Microbiomes 9, (2023).

106. Müller, S. et al. Bacillaene and sporulation protect bacillus subtilis from predation by myxococcus xanthus. Appl Environ Microbiol 80, 5603–5610 (2014).

107. Jumper, J. et al. Highly accurate protein structure prediction with AlphaFold. Nature 596, 583–589 (2021).

108. Grosdidier, A., Zoete, V. & Michielin, O. SwissDock, a protein-small molecule docking web service based on EADock DSS. Nucleic Acids Res 39, 270–277 (2011).

109. Aurelien Grosdidier, Vincent Zoete, O. M. EADock: Docking of Small Molecules Into Protein Active Sites With a Multiobjective Evolutionary Optimization. PROTEINS: Structure, Function, and Bioinformatics 67, 1010–1025 (2007).

